# Temporal single cell atlas of non-neuronal retinal cells reveals dynamic, coordinated multicellular responses to central nervous system injury

**DOI:** 10.1101/2022.07.10.499469

**Authors:** Inbal Benhar, Jiarui Ding, Wenjun Yan, Irene E. Whitney, Anne Jacobi, Malika Sud, Grace Burgin, Karthik Shekhar, Nicholas M. Tran, Chen Wang, Zhigang He, Joshua R. Sanes, Aviv Regev

## Abstract

Non-neuronal cells play key roles in the complex cellular interplay that follows central nervous system (CNS) insult. To understand this interplay at a tissue level, we generated a single-cell atlas of immune, glial and retinal pigment epithelial cells from adult mouse retina before and at multiple time points after axonal transection (optic nerve crush; ONC), identifying rare and undescribed subsets, and delineating changes in cell composition, expression programs, and interactions. Computational analysis charted an inflammatory cascade after injury with three phases. The early phase consisted of reactivation of retinal macroglia and microglia, providing chemotactic signals for immune infiltration, concurrent with infiltration of CCR2^+^ monocytes from the circulation. In the second phase, these differentiated to macrophage subsets resembling resident border-associated macrophages. In parallel, a multicellular interferon program, likely driven by microglia-derived type-I interferon, was synchronously activated across resident glia, expanding beyond rare interferon-responding subsets of glia unexpectedly present in the naïve retina. Our findings provide insights regarding post-injury CNS tissue dynamics and a framework to decipher cellular circuitry, spatial relationships and molecular interactions following tissue injury.

## Introduction

Tissues vary dramatically in their ability to heal and regenerate after sustaining insult. Following injury, cell states, compositions and interactions are dynamically remodeled, thereby affecting the course of healing in both beneficial and detrimental ways (Adler et al., 2020; Meizlish et al., 2021). In particular, the adult mammalian central nervous system (CNS) undergoes extensive remodeling after insult, yet generally fails to regenerate. Immune cells and glia are key orchestrators of tissue repair, but their spontaneous responses to CNS insult are often insufficient to achieve functional recovery, and at times are even counterproductive (Burda and Sofroniew, 2014; Gadani et al., 2015; Schwartz et al., 2020; Shechter and Schwartz, 2013). While much insight has been previously gleaned from studying each cellular component in isolation, a deeper understanding of multicellular networks is required to better understand the mechanisms underlying CNS pathology (Andries et al., 2020; Burda and Sofroniew, 2014; Greenhalgh et al., 2020; Matejuk and Ransohoff, 2020).

With its relative accessibility and highly organized structure, the retina is an invaluable model to study the CNS in general, and its response to injury in particular (Dowling, 2012; London et al., 2013). Optic nerve crush (ONC) injury is a well-established model for studying both neuron-intrinsic and extrinsic processes of CNS degeneration and regeneration (Aguayo et al., 1991; Williams et al., 2020; Yoles and Schwartz, 1998). In this model, transecting the axons of retinal ganglion cells (RGCs), the projection neurons of the retina, results in degeneration of all distal segments and death of ∼80% of RGCs within ∼2 weeks; few if any surviving RGCs regenerate new axons so no visual function is recovered (Williams et al., 2020).

Recent advances in analyzing tissues at single cell resolution are significantly advancing our understanding of tissue biology in homeostasis, injury and disease. We recently used single cell RNA-seq (scRNA-seq) of RGCs during ONC in the mouse to characterize 46 RGC subtypes, based on morphological and molecular features (Tran et al., 2019). We demonstrated dramatic differences in their ability to survive ONC (Bray et al., 2019; Duan et al., 2015; Tran et al., 2019), and highlighted neuron-intrinsic programs that may underlie the selective resilience or vulnerability and regeneration potential of particular RGC subtypes and subclasses (Jacobi et al., 2022; Tran et al., 2019). However, other retinal cells, specifically glia, epithelial and immune cells can impact RGC survival, for example due to benefits from immune-based enhancement of neuroprotection and regeneration (Fisher et al., 2001; Kurimoto et al., 2013; Moalem et al., 1999; Sas et al., 2020; Walsh et al., 2015). Deeper characterization of the heterogeneity of non-neuronal retinal cells at baseline and after ONC is needed to better harness the potential of these cells in healing.

Here, we used the ONC model to study post-injury tissue-level dynamics of non-neuronal cells in the retina of adult mice. We profiled retinal immune, glia and epithelial cells by scRNA-seq in uninjured retina and at six time points from 12 hours to two weeks after ONC. We characterized the heterogeneity of resident cells at baseline and charted the states they acquire after injury. We identified inflammatory cascades whereby early glial reactivation coupled with chemokine upregulation may drive infiltration of circulating CCR2^+^ monocytes after injury, which in turn gives rise to macrophages with overlapping signatures to resident macrophages. In parallel, after injury, an interferon program was synchronously augmented across multiple cell types in a concerted manner, expanding interferon-responsive subsets of retinal glia and microglial cells that surprisingly existed in the naïve (uninjured) retina. In microglia, the interferon state may be a part of their transition between proliferation and maturation. Our findings provide mechanistic insights and a resource for generating and testing further hypotheses in the retina and the CNS at large.

## Results

### A single cell atlas of the retina tissue ecosystem following ONC

To complement our previously profiled atlas of RGCs in uninjured (control; ctrl) adult mouse retina and following ONC (Jacobi et al., 2022; Tran et al., 2019), we generated corresponding scRNA-seq atlases of non-neuronal cells of the retina and the retinal pigment epithelium (RPE) (**Fig. 1a**). From the retina, we used FACS to enrich retinal immune cells, Müller glia and astrocytes (**Methods**) from uninjured (control) retina and at six time points following ONC, through two weeks after injury, when RGC death is at ∼80% (Levkovitch-Verbin et al., 2000; Tran et al., 2019), followed by scRNA-seq (**Fig. 1a**). We complemented this with a cell atlas of the eyecup from uninjured retina and at 12h, 1d, and 2d after ONC (**Fig. 1a**). The RPE is an epithelial monolayer located between the outer retina and the choroid, serving as the outer blood-retinal barrier (oBRB), an immunoregulatory interface between the retina and the circulation, and a site of immune activity after injury (Benhar et al., 2016; McMenamin et al., 2019; O’Koren et al., 2019; Omri et al., 2011).

**Fig. 1:**
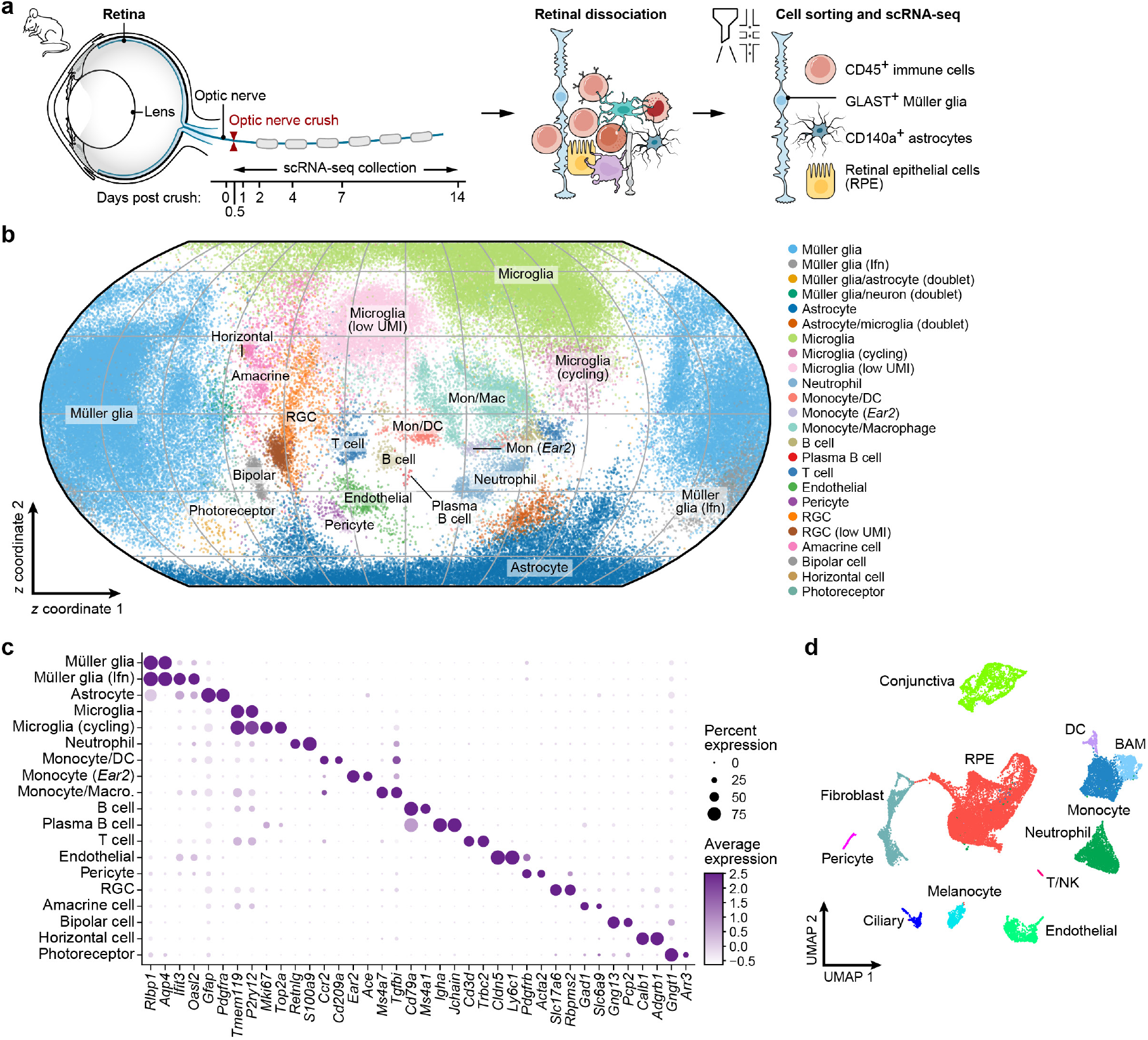
A single cell atlas of the retina tissue ecosystem following ONC. **a,** Experimental overview. **b,c,** Key retinal cell subsets during ONC. **b,** ScPhere embedding of 121,309 single cell profiles (dots) from the retina, projected to 2D by the Equal Earth map projection method, colored by cell type (legend). **c,** Fraction of expressing cells (dot size) and mean expression in expressing cells (dot color) of selected marker genes (columns) across 19 identified retinal cell types, including residual neurons (rows). **d,** 2D Uniform Manifold Approximation and Projection (UMAP) for 21,275 single cells profiled from mouse posterior eyecup at baseline and at 12h, 1d and 2d after ONC, colored by cell type.

From the retina, we recovered 121,309 high quality single cell profiles, annotated them to subsets (putative cell types or states) based on differentially expressed marker genes (**Fig. 1b,c**), identified 107,067 non-neuronal cells, and analyzed them in detail. We first used scPhere (Ding and Regev, 2021), a deep generative model, to integrate the data when introducing time as a batch variable, emphasizing profile similarities across time points (**Fig. 1b** and **Supp. Fig. 1a,b**), to allow effective annotation of cell subsets ahead of comparisons of cells of the same subset across time points (**Methods**). While cell type composition varied along the time course (below), marker gene expression remained sufficiently invariant to annotate 14 major cell types (**Supp. Fig. 1c**).

From the eyecup, we profiled 21,275 cells, and partitioned them into 12 subsets, annotated by expression signatures and marker genes (**Fig. 1d** and **Supp. Fig. 1d,e**) (Geisert et al., 2009; Lehmann et al., 2020; Youkilis and Bassnett, 2021). RPE and immune cells, which we enriched by sorting, comprised 42.3% and 30% of the profiled cells, respectively; the remainder included conjunctival epithelium, scleral fibroblasts, endothelial cells (including *Kdr^high^*and *Kdr^low^* (**Supp. Fig. 1f**) (Lehmann et al., 2020)), melanocytes, ciliary epithelium and pericytes (**Fig. 1d**).

### A temporal inflammatory cascade of infiltrating leukocytes and expanding resident macrophages post ONC

Immune cell composition (**Fig. 2a,b** and **Supp. Fig. 2a**) was dynamically remodeled post ONC, forming a temporal progression of infiltration, proliferation, and expansion of resident and infiltrating cells, with three distinct, albeit partly overlapping, stages.

**Fig. 2:**
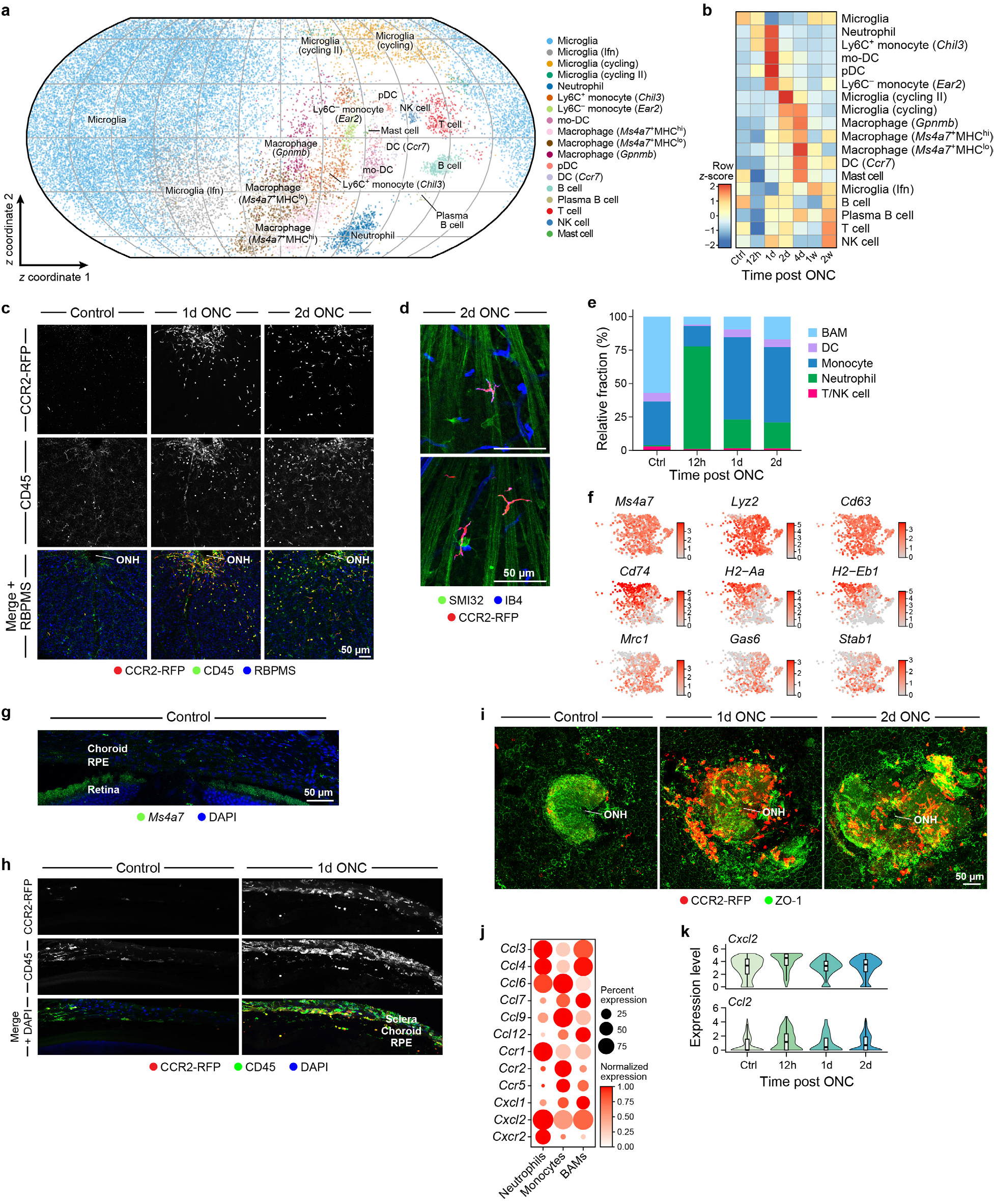
A temporal inflammatory cascade of infiltrating leukocytes and expanding resident macrophages post ONC. **a-b,** Changes in retinal immune cell composition following ONC. **a,** ScPhere embedding of 30,427 immune cell profiles (dots) from the retina, projected to 2D by the Equal Earth map projection method, colored by cell type (legend). **b,** Row z-scores showing the changes in relative abundance of each immune cell subset across time. **c-d,** Blood-derived monocytes infiltrate the retina after ONC. Representative images of IHC on retinal whole-mounts from CCR2^RFP/+^ mice showing CCR2^+^ monocytes (RFP, red) infiltrating the retina and spreading across the GCL (RPBMS, blue) 1-2dpc (**c**), where they come in association with RGC somas and axons (SMI32, green) (**d**). GS-IB4 isolectin (blue) marks blood vessels and activated monocytes/macrophages (n = 2-3 per time point per experiment, representative of two independent experiments). **e-k,** Changes in immune cell composition in the eyecup after ONC. **e,** Fraction (y axis) of immune cell subsets isolated from the eyecup across the time course (x axis). **f,g,** BAMs in the outer eye. **f,** BAMs in the eyecup colored by pan-BAM genes (top), MHC-II^hi^ (middle) and MHC-II^lo^ (bottom) BAM genes (Van Hove et al., 2019). **g,** Representative image of smFISH for *Ms4a7* (green) in naïve (Control) mouse eyecup (n = 3). **h,i,** Monocyte infiltration in the eyecup after ONC. Shown are representative images of IHC on eyecup sections (**h**) or whole-mounts (**i**) from CCR2^RFP/+^ mice, showing sparse CCR2-RFP^+^ cells (red) in uninjured (Control) eyecup, and their increase after ONC, especially around the optic nerve head (ONH) (n = 2 per time point, representative of three independent experiments). **j,k,** Increased chemokine expression by BAMs after ONC. **j,** Fraction of expressing cells (dot size) and normalized expression level (dot color) of chemokines and their receptors by BAMs, neutrophils and monocytes in the eyecup. **k,** Distribution of expression of *Cxcl2* and *Ccl2* by eyecup BAMs across time. Boxplots denote the medians and the interquartile ranges (IQRs). The whiskers of a boxplot are the lowest datum still within 1.5 IQR of the lower quartile and the highest datum still within 1.5 IQR of the upper quartile.

The first stage (12h-1d post crush; pc) was distinguished by substantial increases in neutrophils, Ly6C^+^ monocytes and dendritic cell (DCs). Neutrophils (2.8% of immune cells) were the earliest responders, infiltrating by 12hpc and returning to control levels by 2dpc (**Fig. 2b** and **Supp. Fig. 2b**), and were closely followed by monocytes and DCs, with a more protracted decline. Among the infiltrating monocytes, Ly6C^+^ monocytes (expressing *Ccr2*, *Ly6c2*, *Chil3* and *Plac8*) were the most abundant, rising from 0.4% of immune cells in control retina to 8.6% by 1dpc (Fisher’s exact test, *p* < 0.001). When we tracked monocytes in retinal whole-mounts from CCR2^RFP/+^ transgenic mice (**Fig. 2c**), they were initially concentrated around the optic nerve head (ONH), then spread out peripherally across the nerve fiber, ganglion cell and inner plexiform layers (NFL, GCL and IPL) of the retina, and acquired an elongated morphology indicative of their maturation into macrophages (**Fig. 2c** and **Supp. Fig. 2c**). Some of the CCR2-RFP^+^ cells were in close apposition to RGCs and their axons (**Fig. 2d**). These were followed by a minor subset of Ly6C^-^monocytes (0.7% of immune cells; expressing *Ear2*, *Nr4a1* and *Treml4*, but not *Ccr2* or *Ly6c2* (Giladi et al., 2020; Hanna et al., 2011; Yona et al., 2013)) (**Fig. 2b**). In addition, DCs at this 12h-1dpc window were predominantly from a *Ccr2*^+^ subset of monocyte-derived dendritic cells (moDCs, 1.2% of immune cells; expressing *Ccl5*, *CD209a*, and high levels of MHC-II genes), and a smaller *Ccr9*^+^ plasmacytoid DC subset (pDC, 0.25% of immune cells).

The second phase (2-4dpc) was characterized by microglial proliferation and expansion of macrophage populations (below), as well as rare populations of *Ccr7*-expressing DCs and mast cells (0.2% and 0.04% of immune cells, respectively) (**Fig. 2b**). Finally, in the last phase, at 2wpc, adaptive immune cells, including B, plasma B, T and NK cells, were at their highest, although B, T and NK cells were also present, albeit at lower proportions, in the uninjured retina (**Fig. 2b**). These cells have mostly been studied in the CNS in pathological contexts, but have recently been identified in the naïve brain (Korin et al., 2017; Mrdjen et al., 2018).

Thus, the cellular aspect of the response to CNS injury mirrors the classical wound healing response in peripheral tissues (Gadani et al., 2015; Shechter and Schwartz, 2013), while also providing higher granularity. Importantly, however, unlike wound healing, RGCs generally fail to regenerate. We thus turned to identify cellular and inter-cellular events in the retina and eyecup that may underlie and drive this cascade and its impact on RGCs.

### Immune cell infiltration at the outer blood-retinal barrier early after ONC

We first hypothesized that the early increase in immune cells may be due to infiltration through the oBRB, and thus analyzed 6,415 immune cells from the eyecup atlas in controls and ONC.

Border-associated macrophages (BAMs) were the predominant immune cell subset in the uninjured eyecup (**Fig. 2e**), and expressed macrophage markers and MHC-II genes, as well as *Mrc1*, *Pf4*, *Ms4a7* and *Stab1*, with a range of expression of genes associated with MHC-II^hi^ and MHC-II^lo^ subsets (Jordão et al., 2019; Van Hove et al., 2019) (**Fig. 2f**), similar to macrophages in other CNS compartments (Chakarov et al., 2019; Dick et al., 2022; Eraslan et al., 2022). We validated the presence of *Ms4a7*^+^ and of MHC-II^+^(IA-IE)CD206^+^(Mrc1) cells in the uninjured eyecup (**Fig. 2g** and **Supp. Fig. 2d**).

ONC resulted in dramatic remodeling of the eyecup immune cell composition preceding and mirroring those in the retina (**Fig. 2e**): a transient increase in neutrophils at 12hpc, and a proportional reduction in BAMs, followed by an increase in CCR2^+^ monocytes at 1d and 2d post crush (**Fig. 2e,h**). IHC for monocytes in wholemount eyecups from CCR2-RFP mice indicated that in the control eye, most RFP^+^ cells were posterior to the RPE (**Supp. Fig. 2e**), likely representing choroidal short-lived monocyte-derived macrophages (Joly et al., 2009; O’Koren et al., 2019). Following ONC, more RFP^+^ cells were associated with the RPE (Benhar et al., 2016) (**Fig. 2h**), mostly concentrated around the ONH (**Fig. 2i**). Together with center-to-periphery monocyte spreading across the inner retina (**Fig. 2c**), these results indicate the ONH as the predominant route of leukocyte infiltration into the retina in this model. Notably, receptor-ligand interactions enriched (**Methods**) between BAMs, neutrophils and monocytes involved chemokines and their receptors (**Fig. 2j,k**), indicating the potential role of eyecup resident macrophages in peripheral immune infiltration.

### Early reactivation of retinal glia and microglia

To gain insight into the cues within the retina that could drive temporal progression of the inflammatory response, we analyzed microglia, the CNS-resident macrophages (∼0.5% of retinal cells) and the resident retinal glia (Vecino et al., 2016): Müller glia (∼3% of retinal cells (Jeon et al., 1998; Macosko et al., 2015), expressing *Rlbp1*, *Aqp4* and *Glul;* **Supp. Fig. 3a, top**), which span the entire thickness of the retina and come in contact with all neuronal layers; and astrocytes (∼0.2% of retinal cells, expressing *Pdgfra*, *S100b* and *Gfap,* **Supp. Fig. 3a, bottom**), which reside in the GCL and NFL, in tight contact with the inner retina blood vessels.

Microglia state changes in response to ONC (**Fig. 3a**) were consistent with those to other CNS insults (Hammond et al., 2019; Keren-Shaul et al., 2017; Krasemann et al., 2017; Li et al., 2019; O’Koren et al., 2019; Ronning et al., 2019; Wieghofer et al., 2021), including decreased expression of signature genes as early as 12hpc (**Supp. Fig. 3b**), and increased proliferation 2-4dpc (**Fig. 3a,b**) (Wohl et al., 2009). *In situ* analysis of cycling (KI67^+^) microglia showed their localization in the GCL and in the inner and outer plexiform layers (IPL and OPL) (**Fig. 3c** and **Supp. Fig. 3c,d**).

**Fig. 3:**
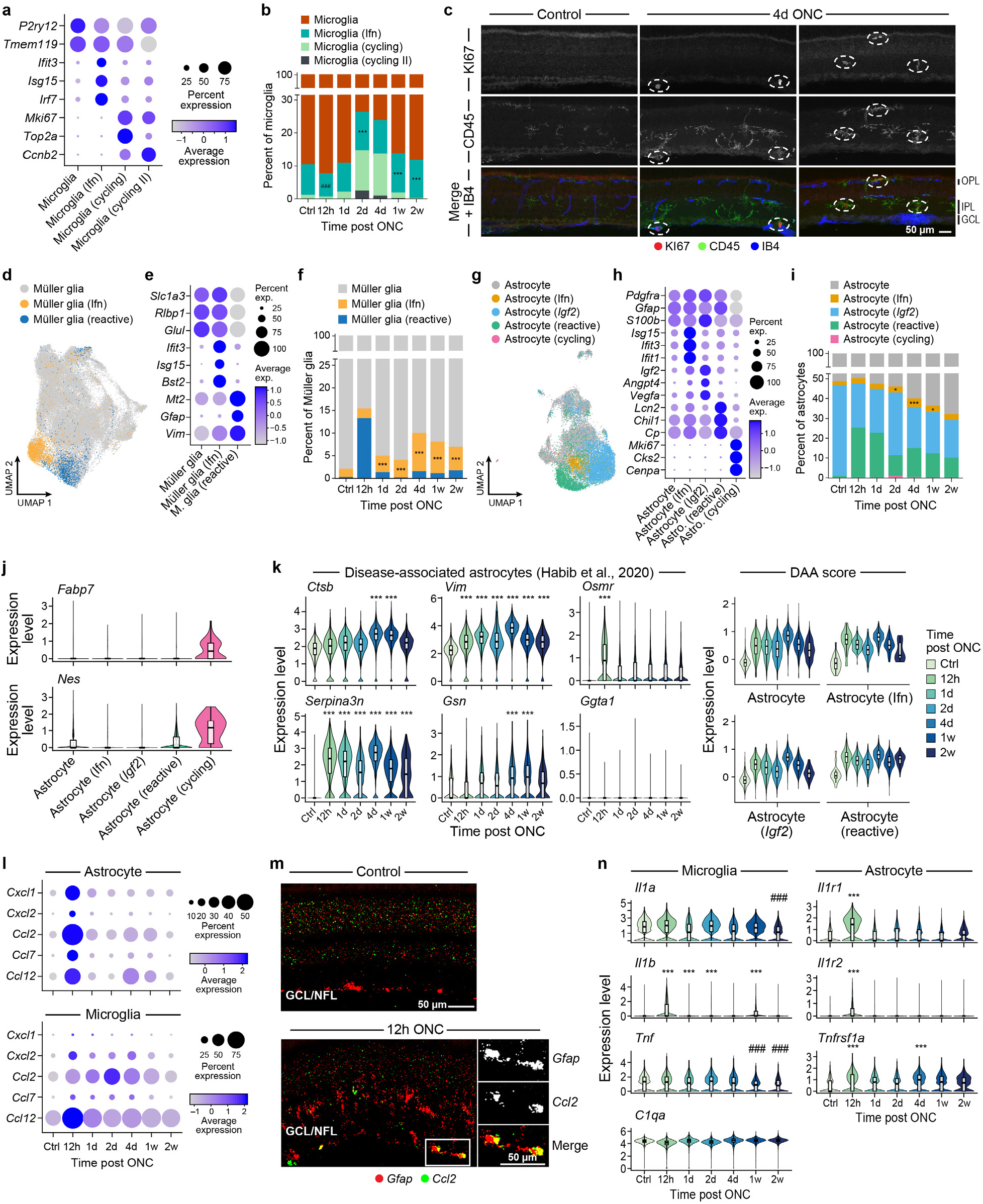
Early reactivation of retinal glia and microglia following ONC. **a,b,** State changes in microglia after ONC. **a,** Fraction of cells (dot size) in each microglia subset, and mean expression level in expressing cells (dot color) of genes differentially expressed between the subsets. **b,** Fraction (y axis) of microglia subsets at each time point (x axis). Asterisks (hashes): time points where the Ifn microglia subset is increased (decreased) relative to ctrl, ***/^###^ *p* < 0.001. **c,** Representative images of IHC on retinal sections showing KI67^+^ (red) cycling microglia (CD45, green) across the retinal layers at 4dpc (n = 3 per time point). **d-k,** Müller glia and astrocyte reactivation following ONC. **d,g,** 2D Uniform Manifold Approximation and Projection (UMAP) for (**d**) 54,565 Müller glia (MG) and (**g**) 18,959 astrocyte cell profiles from control retina and six time points after ONC, colored by cell subset. **e,h,** Fraction of cells (dot size) in each Müller glia (**e**) and astrocyte (**h**) subset, and mean expression level in expressing cells (dot color) of genes differentially expressed between the subsets. **f,i,** Fraction (y axis) of Müller glia (**f**) or astrocyte (**i**) subsets at each time point (x axis). Asterisks: time points where the Ifn MG (**f**) or Ifn astrocyte (**i**) subsets are increased relative to ctrl, \**p* < 0.05, \*\*\**p* < 0.001. **j,** Cycling astrocytes express neural progenitor markers. Distribution of *Fabp7* and *Nes* expression levels (y axis) across astrocyte subsets (x axis). **k,** Retinal astrocytes express disease-associated astrocyte (DAA) genes after ONC. Distribution of expression of signature DAA genes (Habib et al., 2020) across time in astrocytes (left), and as a signature score across astrocyte subsets (right). **l-n,** Glial reactivation and microglial activation provide early immune infiltration signals. **l,** Fraction of expressing cells (dot size) and mean expression level in expressing cells (dot color) of chemokine genes (rows) by astrocytes (top) and microglia (bottom) across time (columns). **m,** Astrocytic expression of the monocyte chemoattractant, *Ccl2*. Representative image of smFISH on retinal sections for *Ccl2* (green) and *Gfap* (red) at 12hpc. The injury-induced increase in *Gfap* expression among Müller glia is also evident (n = 3). **n,** Distribution of expression across time of “A1”-inducing cytokine genes by microglia (left), and of their receptors by astrocytes (right). \*\*\**p* < 0.001, fold change > 1.5 over ctrl; ###*p* < 0.001, downregulated > 1.5-fold.

Müller glia profiles from control and injured retina formed an expression continuum, with pronounced signatures of reactivation early post injury (**Fig. 3d,e**; **Methods**). Reactive Müller glia (2.5% of all Müller glia) were most prevalent at 12hpc (13.3% vs. 0.26% in ctrl; Fisher’s exact test, *p* < 0.001; **Fig. 3f**) with enriched expression of genes associated with reactivation, consistent with the pronounced morphological and expression changes that glial cells undergo following perturbation (Vecino et al., 2016). These included upregulation of *Gfap*, *Vim*, *Mt2*, and decreased expression of Müller glia marker genes (Hoang et al., 2020; Lewis and Fisher, 2003; Vecino et al., 2016) (**Fig. 3e**). Reactive glia also had highest expression of genes induced in mouse Müller glia by excitotoxins and light injury (Hoang et al., 2020) (**Supp. Fig. 3e**), implying common expression programs across injury models.

Similarly, the continuum of astrocytes from control and injured mouse retinas (**Fig. 3g**), also included expansion of reactive astrocytes immediately post-crush (**Fig. 3h,i** from 0.9% at baseline to 25.4% reactive astrocytes at 12hpc; Fisher’s exact test, *p* < 0.001) and declined gradually afterwards (**Fig. 3i**), recapitulating expression programs documented in several injury and disease models (Habib et al., 2020; Hasel et al., 2021; Sofroniew, 2020; Zamanian et al., 2012), including ONC and other retinal pathologies (de Hoz et al., 2016; Liddelow et al., 2017; Qu and Jakobs, 2013). All astrocytes, but especially those in the most “reactive” subset, had increased expression of a pan-reactive astrocyte signature (Liddelow et al., 2017; Zamanian et al., 2012) as early as 12hpc post injury and throughout the time course (**Supp. Fig. 3f**). A minor population (∼0.3%) at 2-4dpc expressed cell cycle genes and higher levels of the neural progenitor markers, *Nestin* and *Fabp7* (**Fig. 3h-j**), suggesting that they are newly proliferating astrocytes, consistent with BrdU labeling experiments in optic nerve injury (Panagis et al., 2005; Wohl et al., 2009). In contrast to this expansion, *Igf2* astrocytes (26.1% of all astrocytes) decreased in proportion post-crush (from 45.9% to 22.2% at 12hpc; Fisher’s exact test, *p* < 0.001; **Fig. 3i**) and expressed higher levels of genes encoding the angiogenesis factors, *Igf2*, *Angpt4* and *Vegfa* (**Fig. 3h**), possibly reflecting their interactions with blood vessels. The astrocyte response in ONC, an acute injury, included increased expression of a signature of disease-associated astrocytes (DAAs) from Alzheimer’s disease (Habib et al., 2020), a chronic neurodegeneration (**Fig. 3k**), suggesting shared features in different neuroinflammatory settings (Habib et al., 2020; Hasel et al., 2021; Wheeler et al., 2020).

The reactive gliosis response in both Müller glia and astrocytes included upregulation of genes related to metal metabolism (**Supp. Fig. 3g**, likelihood ratio test, *p* values adjusted based on BH procedure, **Supp. Fig. 3h**). Trace metals catalyze enzymatic reactions key to visual processing, but their levels must be tightly regulated to avoid neurotoxicity (Picard et al., 2020; Ugarte et al., 2013). The elevated expression of chelators and transporters may be neuroprotective by sequestering excessive metals present in the injured retina, and limiting oxidative stress (Chung et al., 2008; Daruich et al., 2019; Levin and Geszvain, 1998; Li et al., 2017; Picard et al., 2020; Roesch et al., 2012; Ryan et al., 2018; Shu and Dunaief, 2018; Ugarte et al., 2013; Wunderlich et al., 2010).

Taken together, reactivation of both Müller glia and astrocytes is an early event after injury, beginning before substantial changes in RGCs are observed (Tran et al., 2019).

### Glial reactivation and microglial activation provide early immune infiltration signals

The early (12hpc) reactivation of astrocytes and microglia induced cytokines that mediate immune cell recruitment into the CNS in neuroinflammatory conditions (Babcock et al., 2003; Glabinski et al., 1996; Kim et al., 2014; Tanuma et al., 2006). Among these, neutrophil and monocyte chemoattractant expression was upregulated at 12hpc (**Fig. 3l**), coinciding with the emergence of neutrophils and monocytes in the retina (**Fig. 2b**). We confirmed *Ccl2* expression by astrocytes at 12hpc by single-molecule fluorescence *in situ* hybridization (smFISH) (**Fig. 3m**). Thus, astrocytes and microglia activated early after injury may connect injured RGCs and peripheral immune cells, to facilitate the ensuing immune response.

In parallel, microglia-to-astrocyte signaling may promote both neuroprotective and neurotoxic events. In particular, microglial-derived IL1α, TNF and C1q induce a neurotoxic astrocyte state (Liddelow et al., 2017), whereas the IL1β-IL1R1 axis can be neuroprotective (Todd et al., 2019). While *Il1a*, *Tnf* and *C1qa* were constitutively expressed in microglia, *Il1β* was upregulated in microglia at 12hpc, concurrent with elevated expression of genes encoding the IL1 and TNF receptors in astrocytes (**Fig. 3n**). This suggests that induction of neurotoxic astrocytes is an early event, and that neuroprotective and neurotoxic states can co-occur within a cell population and rely on shared signaling pathways.

### A unique retinal pigment epithelial (RPE) cell state emerges early after ONC

To identify additional early events we also partitioned RPE cells into five subsets (**Fig. 4a,b**): four were present at similar frequencies in control and ONC conditions, but one, C12, was highly enriched after ONC (**Fig. 4c**; 2% of RPE cells in control *vs.* 12.1% at 12hpc, 16.8% at 1dpc and 7.12% 2dpc). C12 cells had high expression of genes associated with injury and cell stress, including *Tnfrsf12a*, *Serpina3n* and *Gpnmb*, as well as epithelial-to-mesenchymal transition (EMT) genes, such as *Vim*, *Acta2* and galectins *Lgals1* and *Lgals3* (**Fig. 4d**). We validated the upregulation of *Tnfrsf12a* and Vimentin in RPE after crush (**Fig. 4e,f**). Thus, ONC induced a specific RPE cell state.

**Fig. 4:**
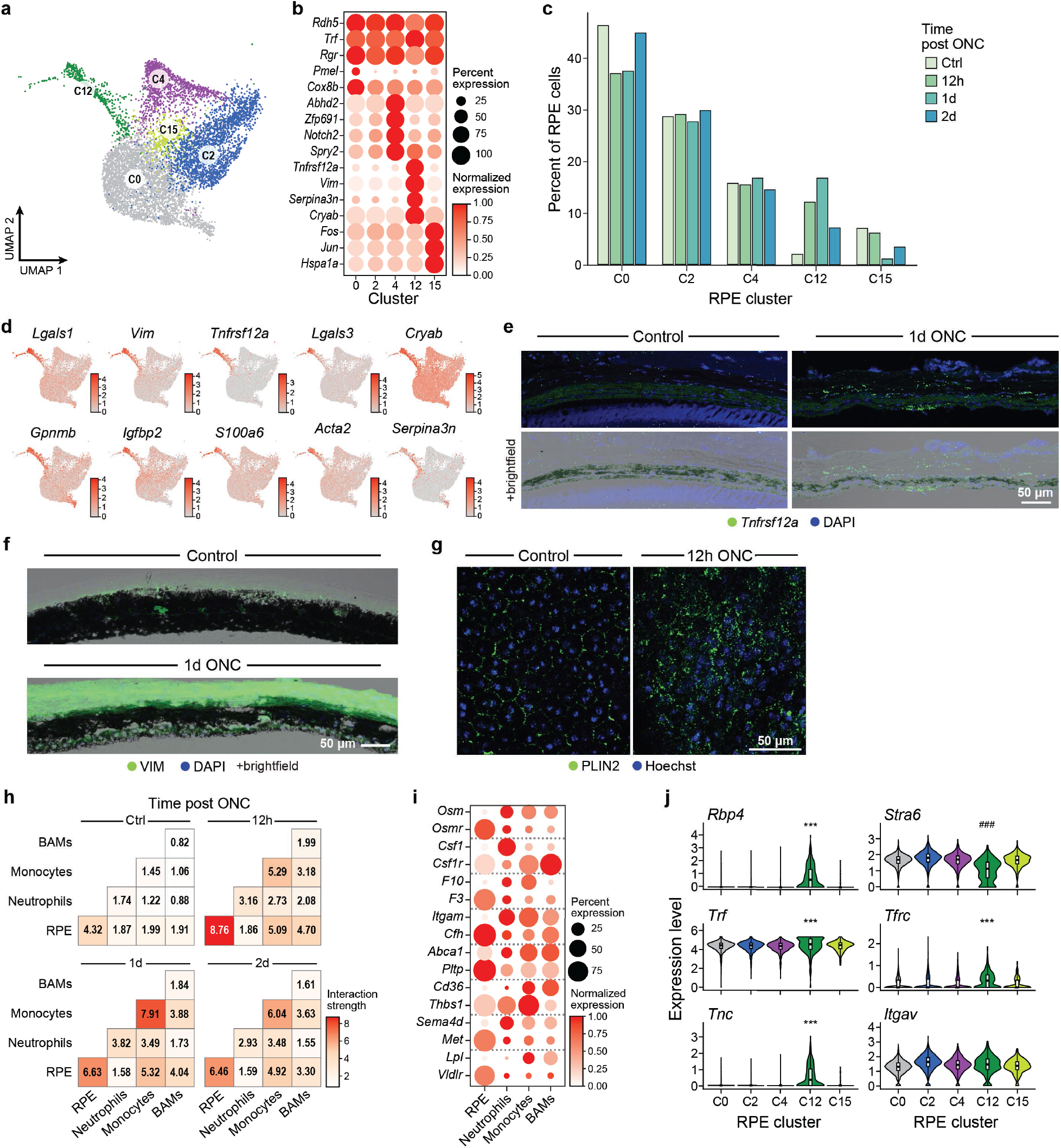
A unique retinal pigment epithelial (RPE) cell state emerges early after ONC. **a-d,** A unique RPE state emerges after ONC. **a,** 2D Uniform Manifold Approximation and Projection (UMAP) for 8,013 RPE cells, colored by subset. **b-c,** RPE heterogeneity. **b,** Fraction of expressing cells (dot size) and normalized expression in expressing cells (dot color) of differentially expressed genes (rows) between RPE subsets (columns). **c,** RPE cluster proportion changes. Shown are cell frequencies (y axis) of RPE clusters (x axis) out of all RPE cells at each time point. **d,** RPE cells colored by expression of C12 top differentially-expressed genes. **e-g,** Expression changes in the RPE after ONC. Representative images of smFISH for *Tnfrsf12a* (green) (**e**), and of IHC for VIM (**f**) and PLIN2 (**g**), showing increased expression after ONC (n = 3 per time point). **h-j,** Interactions between RPE and immune cells increase after ONC. **h,** Changes in interaction strength between pairs of cells from the eyecup across time. **i,** Fraction of expressing cells (dot size) and normalized expression level (dot color) of ligands and their cognate receptors by RPE, neutrophils, monocytes and BAMs in the eyecup. Shown are interactions enriched at 12h-1dpc. **j,** Unique interactions between RPE C12 and the other RPE cells. Shown are distributions of expression of genes encoding ligands and their cognate receptor by RPE subsets.

In addition to the increase in C12 cells, some gene expression changes after ONC were shared in all RPE cells (**Supp. Fig. 4a**). Some of those genes were higher in C12 cells, including *Plin2* (ADRP), a lipid droplet binding protein, which has roles in the visual cycle and in innate immunity (Bosch et al., 2020; Imanishi et al., 2004) (**Supp. Fig. 4b**), the elevation of which we validated *in situ* (**Fig. 4g**). In addition, at 2dpc, there was an increase in the expression of *Nog*, encoding the BMP antagonist Noggin, involved in RPE differentiation (Zahabi et al., 2012) (**Supp. Fig. 4a,c,d**). Thus, although not directly injured, changes in the RPE reflected stress-related processes, such as EMT and dedifferentiation (Stern and Temple, 2015; Tamiya et al., 2010; Yang et al., 2018; Zhao et al., 2011).

Notably, another cluster (C4) expressed development-related genes such as *Abhd2*, *Notch2* and *Spry2* (**Fig. 4b, Supp. Fig. 4e**), enriched for “morphogenesis of an epithelium” (*p* = 3.96×10^-6^), “sensory organ development” (*p* = 3.99×10^-6^) and “embryonic morphogenesis” (*p* = 5.43×10^-6^) genes. Immunostaining confirmed heterogeneity in protein expression of NOTCH2 (**Supp. Fig. 4f**) and ABHD2 (**Supp. Fig. 4g**). This expression state is not related to the circadian rhythm (which controls parts of the RPE transcriptome (DeVera and Tosini, 2020)), as ABHD2 expression pattern was similar at distinct points in the circadian cycle (**Supp. Fig. 4h**). As RPE cells are considered terminally differentiated under normal conditions (Lakkaraju et al., 2020; Stern and Temple, 2015), the functional relevance of this physiological heterogeneity remains to be elucidated.

Receptor-ligand (RL) interactions between RPE, BAMs, neutrophils and monocytes in the eyecup, increased following injury, with intra-RPE and intra-monocyte interactions most prominent, peaking at 12hpc and 1dpc, respectively (**Fig. 4h**). Inferred crosstalk between RPE and immune cells suggested an integrated tissue response as early as 12h-1dpc (**Fig. 4i**), for example between eyecup myeloid cells, expressing *Osm*, encoding oncostatin M, a pleiotropic cytokine shown to be neuroprotective in the retina (Yang et al., 2021), and RPE cells, strongly upregulating its receptor, *Osmr* after ONC (**Fig. 4i** and **Supp. Fig. 4a**). Interestingly, of the top 20 RL pairs in RPE at 12h-1dpc, the vast majority (82.8%) were differentially expressed in RPE C12 cells. Of those, RL pairs that were uniquely expressed after ONC included *Rbp4-Stra6*, involved in retinol transport, *Trf-Tfrc*, involved in metal metabolism, and *Tnc-Itgav*, implicated in retinal and optic nerve pathology (Reinhard et al., 2017) (**Fig. 4j**), suggesting a unique role for C12 RPE cells in remodeling the injured RPE microenvironment.

### *Ms4a7^+^* and *Gpnmb^+^* macrophages increase in retina post-crush

The next, intermediate phase of the inflammatory cascade (**Fig. 2b**) was characterized by the expansion of macrophage subsets and cycling microglia. Three subsets of non-microglial macrophages (**Fig. 2a,b** and **5a**) expressing features of peripherally-derived macrophages and BAMs increased after ONC, rising at 2dpc and peaking at 4dpc (**Supp. Fig. 5a**). All three subsets expressed high levels of *Ms4a7*, *Lyz2* and *Apoe* (**Fig. 5a**) and lower levels of a microglia signature (Bennett et al., 2018; Hohsfield et al., 2020; Jordão et al., 2019; O’Koren et al., 2019; Shemer et al., 2018; Van Hove et al., 2019). *Ms4a7*^+^MHC^hi^ and *Ms4a7*^+^MHC^lo^ subsets were distinguishable by MHC-II expression (**Fig. 5a**), mirroring MHC-II^hi^ and MHC-II^lo^ BAM subsets in homeostatic brain and spinal cord (Jordão et al., 2019; Van Hove et al., 2019) (**Supp. Fig. 5b**). A third subset, *Gpnmb* macrophages, expressed high levels of *Gpnmb*, *Fabp5*, *Spp1* and *Lgals3* (**Fig. 5a**). We confirmed the presence of *Gpnmb^+^* immune cells (*Ptprc^+^*) expressing *Ms4a7* in the inner retina after ONC by smFISH (**Supp. Fig. 5c**).

**Fig. 5:**
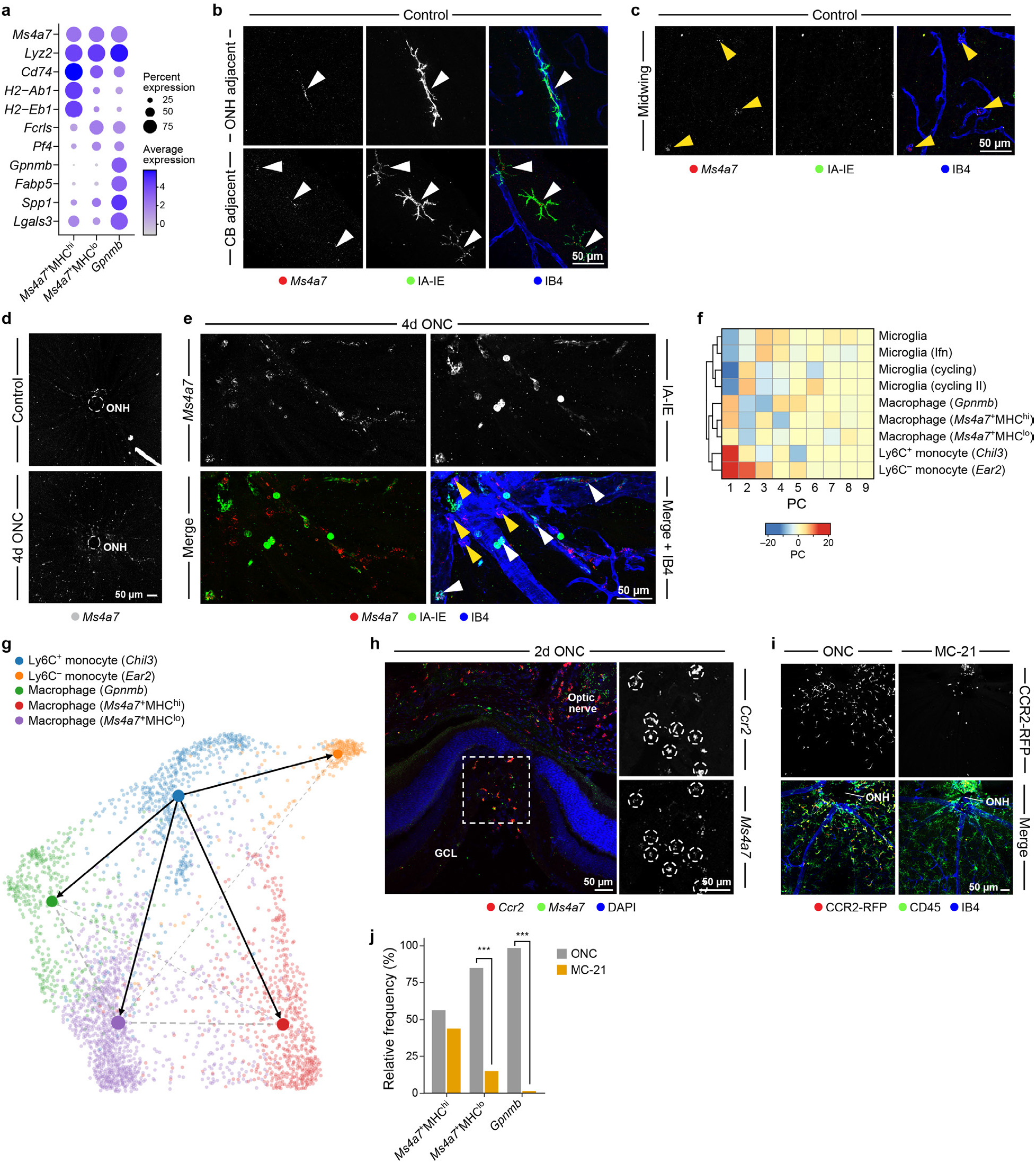
Infiltrating monocytes give rise to macrophage subsets in the injured retina with overlapping expression programs to non-microglial resident retinal macrophages. **a-c,** Three subsets of macrophages identified in the retina. **a,** Fraction of expressing cells (dot size) and mean expression level in expressing cells (dot color) of genes differentially expressed between *Ms4a7*^+^MHC^hi^, *Ms4a7*^+^MHC^lo^ and *Gpnmb* macrophage subsets. **b,c,** Representative images of combined smFISH and IHC on retinal whole-mounts from uninjured (Control) retina, showing *Ms4a7* expression (red) in IA-IE^+^ cells (green), around blood vessels (IB-4, blue), and at the marginal retina, bordering the ciliary body (**b**), and *Ms4a7* (red) expression in IA-IE^-^IB-4^+^ cells (yellow arrowheads) across the GCL (**c**). **d-j,** Infiltrating monocytes give rise to macrophage subsets after ONC. **d,e,** Representative images of smFISH on wholemount retina from uninjured (Control) and 4dpc retina showing increase in *Ms4a7^+^* cells after injury, including *Ms4a7*^+^IA-IE^+^ (white arrowheads) and *Ms4a7*^+^IA-IE^-^ (yellow arrowheads) cells. A zoomed-in inset from the 4dpc image in **d** is shown in **e** (n = 2-3 per time point, representative of three independent experiments). **f,** Cluster analysis showing transcriptional proximity of mononuclear phagocyte subsets in the retina. **g,** Force-directed layout view of monocyte and macrophage subsets with predicted cell transition vectors (arrows) based on partition-based graph abstraction (PAGA), colored by cell type. **h,I,** Representative images of smFISH on 2dpc retinal sections (**h**), showing co-expression of *Ccr2* (red) and *Ms4a7* (green), and of IHC on 2dpc retinal whole-mounts (**i**) for CD45 (green), IB4 (blue) and CCR2-RFP^+^ (red) showing the depletion of monocytes from the retina after MC-21 treatment (n = 3 per condition). **j,** Macrophages are reduced in the retina after monocyte depletion. Relative frequency (y axis) of *Ms4a7* and *Gpnmb* macrophage subsets (x axis) in the retina at 1wpc, with and without MC-21 treatment (pool of n = 5 mice per condition; n = 4633/1551 and 2810/96 single immune cells/macrophages from ONC and MC-21, respectively). \*\*\**p* < 0.001.

Both *Ms4a7*^+^MHC^hi^ and *Ms4a7*^+^MHC^lo^ macrophages were present at lower proportions in uninjured retina (**Supp. Fig. 5a**), and had distinct morphologies and spatial distributions. We found IA-IE^+^ (MHC-II^+^) cells in the naïve retina, wrapped around blood vessels in the juxtapapillary area (ONH), and at the retinal margins adjacent to the ciliary body (CB) (Dick et al., 1995; O’Koren et al., 2016; Xu et al., 2007a) (**Supp. Fig. 5d**), and showed *Ms4a7* expression among these perivascular and CB-adjacent IA-IE^+^ cells (**Fig. 5b**). *Ms4a7^+^*IB-4^+^IA-IE^-^ cells *in situ*, likely corresponding to the *Ms4a7*^+^MHC^lo^ scRNA-seq subset, were morphologically and spatially distinct, dispersed across the inner surface of the retina (**Fig. 5c**). Thus, resident retinal macrophages, expressing *Ms4a7*, include the MHC-II^+^ population and an additional, previously undescribed MHC-II^lo^ subset. The location of *Ms4a7*^+^MHC^lo^ macrophages abutting the vitreous, which separates the retina from the anterior, non-CNS parts of the eye, supports their identity as a distinct, newly-identified subset of retinal BAMs.

### Macrophages in the injured retina arise from infiltrating Ly6C^+^ monocytes and share expression programs with non-microglial resident retinal macrophages

There were more *Ms4a7*^+^ cells around the ONH after ONC (**Fig. 5d**), including both *Ms4a7^+^*IA-IE^+^ and *Ms4a7^+^*IA-IE^-^ cells (**Fig. 5e**). Since *Ms4a7* and *Gpnmb* macrophage subsets were closer in their expression profiles to monocytes than to microglia (**Fig. 5f**) and increased after the infiltration of monocytes (**Fig. 2b**), we hypothesized that they originated from the infiltrating monocytes arising at earlier time points in the eyecup and retina (**Fig. 2b,e**).

Trajectory and RNA velocity analysis (La Manno et al., 2018) (**Fig. 5g**) predicted that Ly6C^+^ monocytes not only gave rise to Ly6C^-^ monocytes (as in fate-mapping studies (Yona et al., 2013)), which are a “sink” not contributing to the macrophage populations (Trzebanski and Jung, 2020; Yona et al., 2013), but also branched into *Ms4a7* and *Gpnmb* macrophages (**Fig. 5g**), supporting our hypothesis that infiltrating monocytes gave rise to these cells, with each as a separate branch. We obtained similar results by an independent analysis using a computational approach that leverages the time point data based on optimal transport (Waddington-OT (Schiebinger et al., 2019)) (**Supp. Fig. 5e**). Moreover, *Ccr2*- and *Ms4a7*-expressing cells overlapped in the retina at 2dpc (**Fig. 5h**).

To experimentally test whether infiltrating monocytes give rise to *Ms4a7* macrophages after ONC, we depleted circulating CCR2^+^ monocytes during the first week post injury using systemic administration of an anti-CCR2 antibody, MC-21 (Giladi et al., 2020; Mack et al., 2001). MC-21 treatment efficiently depleted monocytes in the retina (**Fig. 5i**), and decreased macrophage subsets at 1wpc (by scRNA-seq) (**Fig. 5j**). Thus, these results support the hypothesis that infiltrating CCR2^+^ monocytes contribute to the post-injury increase in *Ms4a7* and *Gpnmb* macrophages.

### Ifn microglia increase in ONC as an intermediate state between microglia proliferation and maturity

In parallel to the increase in monocyte-derived macrophages, as the inflammatory cascade advanced, there was also proliferation of microglia at 2-4dpc (**Fig. 2b**), followed by a moderate increase in a subset of Ifn microglia (at 1-2wpc) that was also present at lower proportions in the control retina (9.3% of microglia) (**Fig. 2b,3b**). Ifn microglia have mainly been associated with aging and neurodegeneration (Deczkowska et al., 2017; Dorman et al., 2021; Grabert et al., 2016; Hammond et al., 2019; Mathys et al., 2017; O’Koren et al., 2019; Roy et al., 2020), and, to our knowledge, were not previously described in the naïve adult CNS. IFIT3^+^ myeloid cells were present in uninjured (control) retina *in situ*, in the IPL and OPL, where microglia primarily reside in uninjured retina, as well as in the GCL (**Fig. 6a**).

**Fig. 6:**
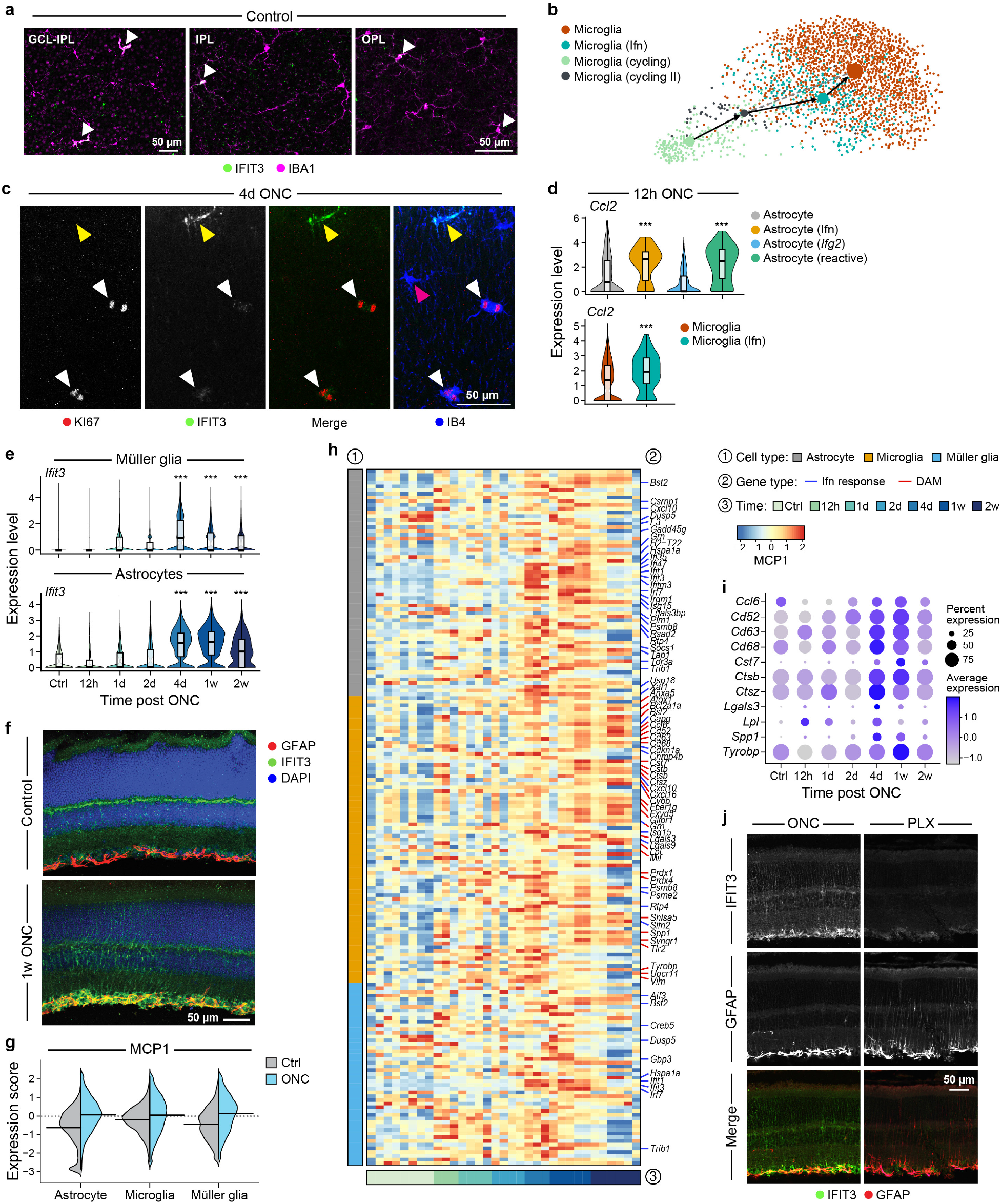
The interferon-response is coordinated in the retina after ONC. **a-c,** Ifn microglia are an intermediate state between microglia proliferation and maturity. **a,c,** Representative images of IHC on retinal whole-mounts showing IFIT3^+^ (green) microglia (IBA1, magenta) (white arrowheads) in the naïve retina, at the levels of the GCL, IPL and OPL (**a**), and KI67^+^IFIT3^+^ microglia (white arrowheads) at 4dpc (**c**). The yellow arrowhead depicts an IFIT3^+^KI67^-^ microglial cell, and the magenta arrowhead depicts an IFIT3^-^KI67^-^ microglial cell (all are IBA1^+^) (n = 3 per time point). **b,** Force-directed layout view of microglia subsets with PAGA vectors, colored by cell subset, showing Ifn microglia at an intermediate stage between cycling and mature microglia. **d,** *Ccl2* expression by astrocytes and microglia at 12hpc is higher in the Ifn subsets. Shown is the distribution of expression of *Ccl2* by astrocyte (top) and microglia (bottom) subsets at 12hpc. \*\*\**p* < 0.001. **e,f,** Rare interferon-responding retinal Müller glia and astrocyte populations expand following ONC. **e,** Distribution of *Ifit3* expression levels (y axis) at each time point (x axis), in Müller glia (top) and astrocytes (bottom). **f,** Representative images of immunohistochemistry (IHC) on sections from control (top) and 1wpc (bottom) retina showing IFIT3 expression (green) on GFAP^+^ (red) astrocytes and Müller glia (n = 3 per time point). Blue: DAPI. **g-i,** A multicellular interferon response program is increased in the retina after ONC. **g,** Distribution of overall expression scores (y axis) of MCP1 across the different cell types (x axis) for cells from uninjured (Ctrl; gray) and ONC (light blue) retinas, showing that MCP1 is induced in ONC samples. **h,** Average expression (z-score, red/blue bar) of upregulated genes (columns) in MCP1, sorted by cell type (left color bar) and across time (bottom color bar). Ifn-related genes are marked with blue tick marks, DAM genes with red. **I,** Fraction of expressing cells (dot size) and mean expression level in expressing cells (dot color) of expression of DAM genes from MCP1 on microglia across time. **j,** Representative images of IHC on 1wpc retinal sections from mice treated with the CSF1R-inhibitor, PLX5622 (right; PLX) and controls (left; ONC), showing decreased IFIT3 (green) and increased GFAP (red) expression after myeloid cell depletion (n = 3 per condition).

RNA velocity analysis (La Manno et al., 2018) of microglia profiles supported a model where cycling microglia transition through the Ifn state as they mature (**Fig. 6b**). Consistent with this model, KI67^+^ cells in the post-crush retina expressed IFIT3 protein, though at lower intensity than in IFIT3^+^KI67^-^ cells (**Fig. 6c**). This model is also in line with reports on microglia repopulation after depletion in the adult brain (Zhan et al., 2019). Indeed, brain-repopulating microglia profiles from a separate dataset (Huang et al., 2018a) expressed significantly higher levels of 37 ISGs that were differentially expressed between our Ifn and control microglia (**Supp. Fig. 6a**). Together, these results suggest that an interferon program is activated during microglia self-renewal in the adult CNS as a transitional state towards maturation.

### Rare interferon-responding retinal Müller glia and astrocyte populations also expand following ONC

Surprisingly, in parallel to Ifn microglia, small subsets of Müller glia (4.1% of all Müller glia) and astrocytes (3.1%) expressing interferon-stimulated genes (ISGs, *e.g.*, *Ifit1*, *Ifit3*, *Isg15*; **Figure 3e,h** and **Supp. Fig. 6b,c**), increased in proportion from 1dpc, peaking 4d-1wpc, although also detected in the retina at all time points, including pre-injury (**Fig. 3f,i**, Fisher’s exact test). ISG expression in retinal glia was reported in autoimmune inflammation (Heng et al., 2019), but to our knowledge, neither after ONC nor in the naïve retina. In the brain, Ifn-responsive astrocytes have been recently identified as a rare subset across CNS pathology models, but, unlike the retina, were virtually absent in the healthy state (Hasel et al., 2021). This suggests that retinal glia maintain populations poised to respond to immune challenge, for instance by promoting infiltration of circulating leukocytes upon need. Notably, *Ccl2* expression at 12hpc was higher in Ifn subsets of astrocytes and microglia compared to their corresponding baseline populations (*p* < 0.001, log2 fold-change = 0.54 for astrocytes; 0.47 for microglia; **Fig. 6d**).

Later after injury, the cell-intrinsic expression of some ISGs, such as *Ifit3*, increased in all astrocytes and in Müller glia (**Fig. 6e, Supp. Fig. 6d**), which we validated at the protein level (**Fig. 6f**), indicating propagation of the Ifn response across the retina. An “A1” astrocyte reactivity signature, acquired by astrocytes in neuroinflammatory settings (Guttenplan et al., 2020; Liddelow et al., 2017; Zamanian et al., 2012), which peaked at 1wpc, was most pronounced in Ifn astrocytes (**Supp. Fig. 6e**).

### The interferon response is coordinated across cell types following injury and may be driven by microglia-produced interferon-beta

Given the increase in each of Ifn Müller glia, Ifn astrocytes, and Ifn microglia post-ONC, we hypothesized that this response may be coordinated across these cell types. To test this hypothesis, we used DIALOGUE (Jerby-Arnon and Regev, 2022), a method that finds gene programs whose expression is coordinated across cells, to identify multicellular programs (MCPs) of genes whose expression is co-regulated in Müller glia, astrocytes and microglia across the 33 samples (**Methods**).

The top MCP higher in ONC *vs*. control (**Fig. 6g, Supp. Fig. 6f**) was an Ifn-response program coordinated at the tissue level (37 genes shared between 149 MCP genes and 878 interferon response genes (from ImmGen), Fisher’s exact test, *p* < 0.001; **Fig. 6h,** blue tick marks on right; **Supp. Table 1**). The MCP peaked around 4 days to 1 week post injury (**Supp. Fig. 6g**). The microglia component of this MCP was enriched for disease-associated microglia (DAM) genes (30 genes shared between 149 MCP genes and 321 DAM signature genes (Keren-Shaul et al., 2017; Krasemann et al., 2017; O’Koren et al., 2019); Fisher’s exact test, *p* < 0.001; **Fig. 6h,** red tick marks on right, **6i**, **Supp. Table 1**). Type I Ifn and DAM programs are correlated and partly overlap in microglia in models of neurodegeneration (Mathys et al., 2017; Roy et al., 2020), though their functional interrelationship is unclear. DAMs were linked to phagocytosis of amyloid plaques (Keren-Shaul et al., 2017) and developmental apoptosis (Anderson et al., 2019), while Ifn induced complement-mediated synaptic pruning (Roy et al., 2020). These programs may be related to the clearance of dying RGCs in ONC.

To identify potential cellular sources of interferon activating the multicellular program, we analyzed the levels of interferon transcripts across the cells and time points in our atlas (**Supp. Fig. 6h**). *Ifnb1* was mainly expressed by microglia subsets, including the Ifn microglia themselves, increasing starting at 2dpc (**Supp. Fig. 6h**). Lower levels of *Ifnb1* expression were also found in astrocytes and Müller glia, RGCs and Ms4a7 macrophages, whereas *Ifng* was mostly expressed by T and NK cells, especially at the 2w time point (**Supp. Fig. 6h**). Thus, a type I interferon (Ifn-I) response induced by microglia is the most likely driver in the time window of rapid RGC loss (Tran et al., 2019) and highest activation of the multicellular interferon program (**Supp. Fig. 6g**). This is in contrast to ISGs activation in the uveitic retina, which was proposed to be due to IFN-gamma from infiltrating Th1 cells (Heng et al., 2019).

We supported our findings by treating mice with PLX5622, a CSF1R inhibitor that effectively depletes retinal microglia (Huang et al., 2018b) and other myeloid cells. Retinas from PLX-treated mice showed diminished expression of IFIT3 and stronger GFAP (**Fig. 6j**), further indicating the involvement of myeloid cells in regulating glial reactivation.

### Multicellular interactions in the injured retina involve glial activity and decreased RGC function

To assess how cell-cell interactions might overall regulate cellular dynamics and especially might impact RGCs in the injured retina, we computed potential RL interactions between all the major resident cells from our retina data, including RGCs (Tran et al., 2019) (**Fig. 7a**). Overall, microglia interacted most strongly with other cell types (**Fig. 7a**), illustrating their constant surveillance of the retinal microenvironment. Early (12h-1dpc) interactions after ONC were dominated by astrocytes, including genes associated with extracellular matrix remodeling, such as *Fn1-Itgav* and *Timp1-Cd63* (**Fig. 7b**). The Il1 axis between microglia and astrocytes (above) was also an enriched interaction at 12hpc. At later time points, interactions involving microglia, astrocytes, and Müller glia reflected immunoregulation and restoration of homeostasis, including through *Grn-Tnfrsf1a* (Kuse et al., 2017), *Serpinf1-Plxdc2* (Vigneswara et al., 2013), *Psap-Gpr37/Gpr37l1* (Jolly et al., 2018) and the TGFβ axis (Ma et al., 2019) (**Fig. 7c**).

**Fig. 7:**
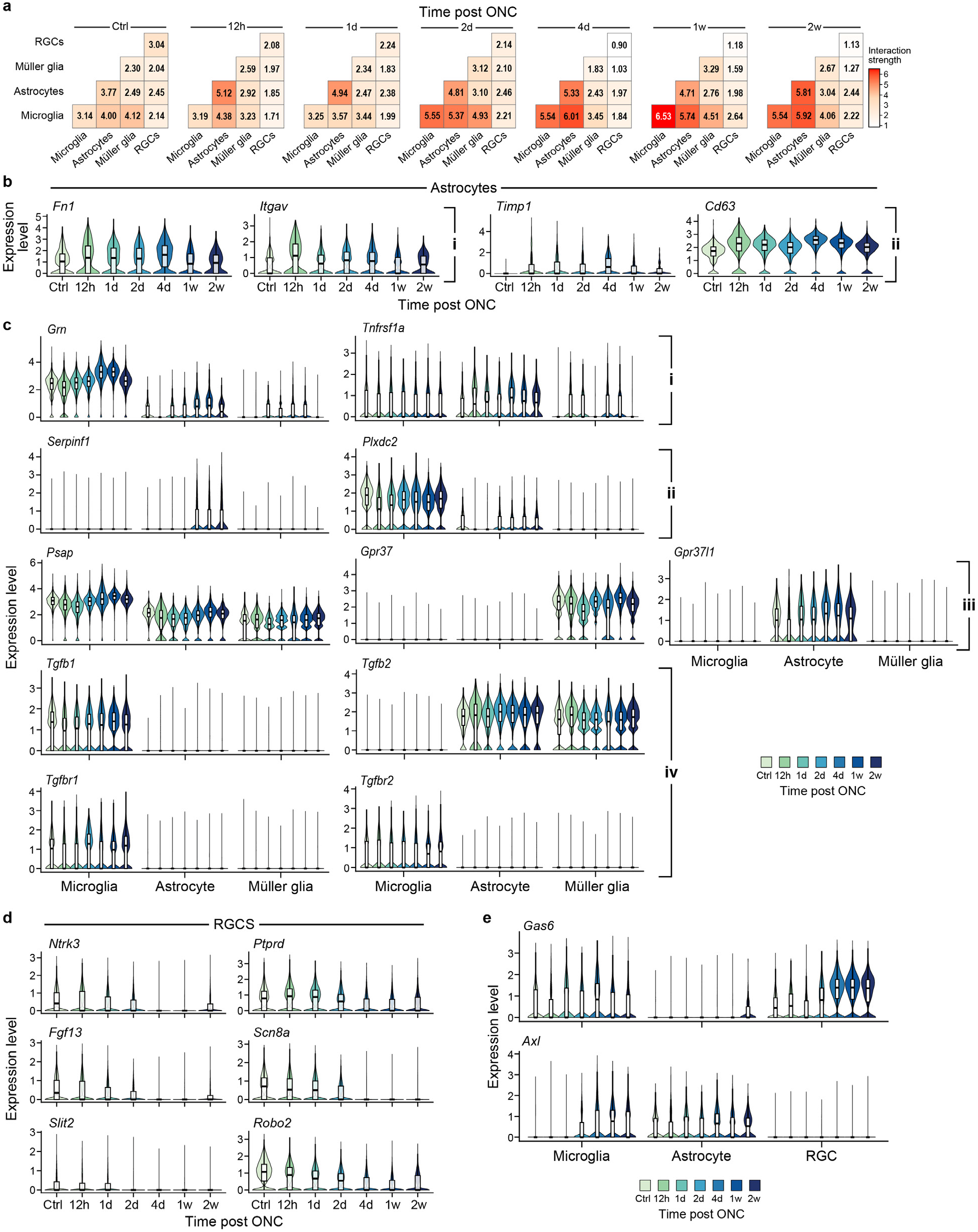
Multicellular interactions in the injured retina involve glial activity and decreased RGC function. **a,** Changes in interaction strength between pairs of retinal cell types across time. **b-e,** Distribution of expression across time of genes encoding receptor and ligand pairs by astrocytes, RGCs, microglia and Müller glia.

Conversely, RGC-RGC interactions became weaker starting at 4dpc (**Fig. 7a**), corresponding to the time of initial reduction in RGC numbers following ONC (Tran et al., 2019). These included interactions involved in neurotrophic signaling (e.g., *Ntrk3*-*Ptprd*), neuronal firing (*Fgf13*-*Scn8a*) and axon guidance (*Slit2-Robo2*) (**Fig. 7d**), reflecting a decline in homeostatic functions after injury (Tran et al., 2019). Similarly, the strength of RGC-Müller glia interactions, including cell-adhesion molecules that modulate neuronal functions (*e.g.*, *Nlgn1-Nrxn2/Nrxn3*, *Sltirk2-Ptprs/Ptprd*), decreased with time (**Fig. 7a**). While the strength of RGC interactions with astrocytes and microglia changed little over time (**Fig. 7a**), RL pairs at the later time points included *Axl-Gas6* (**Fig. 7e**), which have complex roles in the CNS (Fourgeaud et al., 2016; Tondo et al., 2019), and might mediate phagocytic clearance of apoptotic RGCs. Thus, while inter- and intra-glia and immune cell interactions are highly dynamic, those between RGCs and glia or immune cells show little dynamism, such that either RGC death underlies the dearth of interactions, intrinsic events dominate their fate, or the dearth of interactions may even contribute to RGC death.

### Expression of genes associated with human retinal disease across mouse cell types

Finally, we assessed the expression in mouse of genes whose orthologs are associated with human retinal diseases. We included cell types across our entire time course, reasoning that some disease-associated genes reflect impact on disease initiation and others on disease progression.

We first examined glaucoma, a major cause of blindness, characterized by RGC loss (Wang et al., 2021), analyzing genes associated with primary open-angle glaucoma (POAG) (Gharahkhani et al., 2021), and with two optic nerve head parameters associated with glaucoma risk: vertical cup-to-disc ratio (VCDR) and vertical disc diameter (VDD) (Han et al., 2021). Genes associated with POAG, VCDR and VDD (**Supp. Table 2**) were expressed in various cell types, including RGCs, astrocytes and Müller glia, as reported in human (Yan et al., 2020), as well as in the vascular compartment, including endothelial cells, pericytes and smooth muscle cells (**Supp. Fig. 7**). Many of these genes were highly enriched in cycling astrocytes, a minor population in our model. Astrocyte proliferation within the ONH was observed in models of experimental glaucoma, and in human patients at the early phases of the disease (Lozano et al., 2019; Prasanna et al., 2011), likely in response to elevated intraocular pressure, currently the major modifiable risk factor in glaucoma. Notably, we did not observe dependence with time, substantial differential expression, or enrichment in the key cell types associated with the response.

When analyzing genes associated with a variety of diseases in which photoreceptor dysfunction or death leads to blindness (retinitis pigmentosa, congenital stationary night blindness, Leber congenital amaurosis, Usher syndrome, hereditary and age-related macular degeneration), many genes were expressed in the RPE, an important player in the pathogenesis of many retinal diseases. For instance, in retinitis pigmentosa, RPE cells were the main expressers of 14 associated genes, including *Rlbp1*, *Rpe65*, *Abca4*, *Lrat* and *Rgr* (**Supp. Fig. 8**), consistent with reports in human (Orozco et al., 2020). Although mice do not have a macula, some aspects of macular degeneration can be modeled in mice, particularly those that pertain to the RPE (Cheng et al., 2020; Ibbett et al., 2019). Indeed, RPE and additional eyecup cells were enriched for genes associated with macular dystrophies and with age-related macular degeneration, including *Abca4*, *Otx2*, *Fbln5* and the complement factor *Cfh*. Additional complement genes and Toll-like receptor genes were enriched in retinal glia and myeloid cells. This underscores the utility of our atlas in probing expression of disease relevant genes in the mouse.

## Discussion

Here, we provided an atlas of the non-neuronal cells in the retina and charted the tissue response to severe acute CNS injury, delineating changes in non-neuronal retinal cells along a time course corresponding to RGC degeneration (**Fig. 8**). Our analysis shows that an inflammatory cascade was initiated early after injury (phase I), prior to RGC death, with activation of resident glia involving chemokine signals for leukocyte infiltration, which was observed in the oBRB and inner retina. At intermediate time points (phase II), concurrent with the peak rate of RGC death (Tran et al., 2019), infiltrating monocytes differentiated into distinct macrophage subsets, which overlapped in gene expression with resident retinal BAMs. In parallel, during this phase, we identified a synchronous Ifn-response program that was induced in microglia, astrocytes and Müller glia, possibly triggered by DNA damage (Cox et al., 2015; Härtlova et al., 2015; Mathys et al., 2017) as part of the apoptotic process in injured RGCs (Tran et al., 2019). Finally, at 1-2wpc (phase III), there were indicators of restoration of homeostasis, such that glial cell proportions begin to return to their baseline levels (**Fig. 3b,f,i**), with enriched interactions among them including TGFβ signaling.

**Fig. 8:**
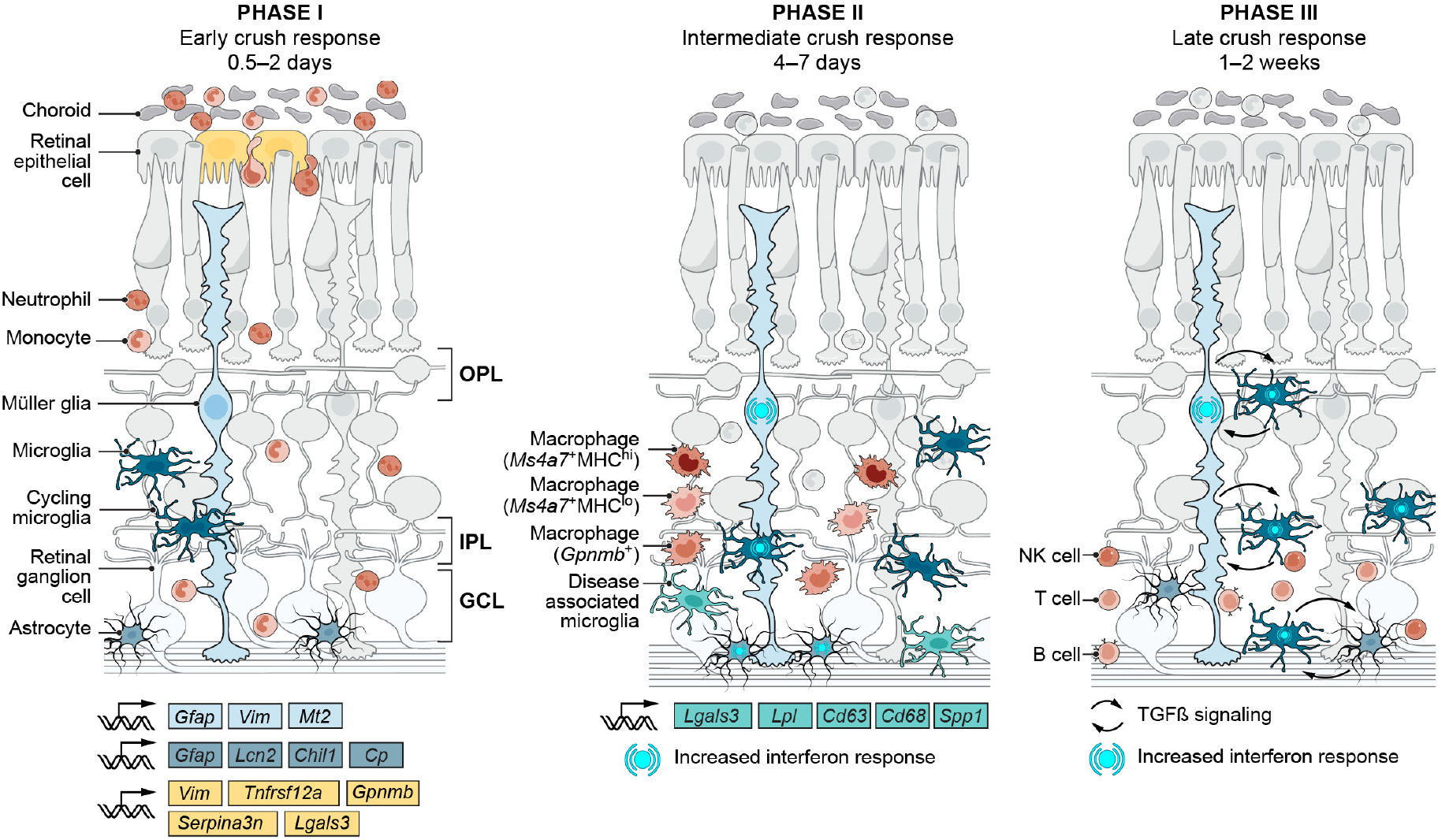
Schematic model summarizing the tissue dynamics along the response to ONC.

The Ifn-I response has multifaceted immunomodulatory roles in the CNS, at both homeostatic and pathological conditions (Abdullah et al., 2018; Baruch et al., 2014; Blank and Prinz, 2017; Deczkowska et al., 2016; Khorooshi and Owens, 2010; Rothhammer et al., 2016; Roy et al., 2020). Ifn-I signaling in the retina is mostly associated with protective immunosuppression (Hooks et al., 2008; Lückoff et al., 2016; Wang et al., 2019), possibly linking the post-injury tissue-wide expansion of the Ifn response with inflammatory resolution at later time points. Notably, we recently found that an Ifn program is also activated in RGCs that regenerate after ONC in response to growth-promoting interventions (Jacobi et al., 2022), further indicating the potential of well-controlled immune responses to benefit CNS repair.

To our surprise, we detected Ifn-responding glial subsets not only after injury, but also in the uninjured retina, prompting the question of whether parallel subsets exist in other CNS compartments under homeostasis, which were perhaps overlooked due to their rarity. Homeostatic Ifn signaling may keep low-grade inflammation in check, with resident cells poised to rapidly respond to perturbation (Goldmann et al., 2015; Lindqvist et al., 2016; Taniguchi and Takaoka, 2001). Supporting this hypothesis, early chemokine expression by astrocytes and microglia was higher in their respective Ifn subsets. Future studies will elucidate the effects of the interferon program in the naïve retina and in the context of ONC, and its consequences on RGC survival and the inflammatory cascade.

In microglia, the Ifn-I program has mainly been associated with aging and neurodegeneration (Deczkowska et al., 2017; Dorman et al., 2021; Grabert et al., 2016; Hammond et al., 2019; Mathys et al., 2017; O’Koren et al., 2019; Roy et al., 2020). Our findings indicate that the Ifn-I response is also activated during microglial renewal in the adult CNS after acute injury and, as found in other studies, in repopulation after CSF1R inhibition (Huang et al., 2018a; Zhan et al., 2019). Whether homeostatic self-renewal of microglia also involves activation of an interferon-response program remains to be determined.

Our findings expand the scope of previously-described ocular macrophage subsets (Jordão et al., 2019; Mrdjen et al., 2018; Van Hove et al., 2019; Wieghofer et al., 2021), as we identified several macrophage populations in the retina distinct from microglia, most prominently *Ms4a7*^+^MHC-II^hi^ and *Ms4a7*^+^MHC-II^lo^ subsets, which comprised both resident and infiltrating macrophages. Intriguingly, the presence of spatially-segregated MHC-II-high and low resident macrophages recurs in health and disease across mouse and human tissues (Chakarov et al., 2019; Dick et al., 2022; Eraslan et al., 2022). In naïve retina, *Ms4a7*^+^MHC^hi^ macrophages were located along the vasculature, and lined the marginal retina at the seam with the ciliary body, which makes up the blood-aqueous barrier and can serve as a monocyte entry site to the retina after injury (Joly et al., 2009; Shechter et al., 2013). *Ms4a7*^+^MHC^lo^ cells, which, to our knowledge, have not been previously described in the retina, were interspersed across its inner surface, bordering with the vitreous body, an immune-privileged region between the lens and retina. We further characterized resident macrophages at the outer blood-retinal barrier, where molecular markers with which to access them were previously lacking (McMenamin et al., 2019). The localization of resident macrophages at border regions in the CNS implicates them as tissue sentinels, sensing and communicating deviation from homeostasis (Kierdorf et al., 2019). Future studies, informed by the markers we identified, can help determine the functional relevance of these macrophage subsets to retinal homeostasis.

Our findings support a model where, in the context of optic nerve injury, CCR2^+^ monocytes give rise to *Ms4a7*^+^ macrophage subsets. This model is based on the increase in macrophages after ONC, our trajectory analyses of cell transitions, the co-expression of *Ccr2* and *Ms4a7* in infiltrating retinal cells, and the reduction in macrophages after monocyte depletion. Whether these cells integrate into the resident macrophage repertoire remains to be determined. Studies in the brain have shown that most BAMs are derived from embryonic erythro-myeloid precursors, are long-lived and self-renewing. However, in adulthood, dural and choroid plexus macrophages are replenished by circulating monocytes (Goldmann et al., 2016; Mrdjen et al., 2018; Utz et al., 2020; Van Hove et al., 2019). Interestingly, a recent report demonstrated that brain perivascular macrophages originate from meningeal macrophages postnatally (Masuda et al., 2022). In the naïve retina, the vast majority of resident macrophages are similarly of embryonic origin and long-lived, though a population of shorter-lived MHC-II^hi^ macrophages has been described (O’Koren et al., 2019; Wieghofer et al., 2021). These distinctions are blurred in our system and in other neuroinflammatory conditions, which involve infiltration of blood-derived myeloid cells, and phenotypic changes among resident macrophages (Ajami et al., 2018; Jordão et al., 2019). Notably, resident macrophages are prone to replacement by blood-derived myeloid cells after injurious perturbations, such as irradiation (Bechmann et al., 2001; Hickey and Kimura, 1988; Kierdorf et al., 2019; Xu et al., 2007b). ONC might similarly invoke replacement of resident macrophages. A combination of fate mapping and spatial genomics can help determine the ontogeny and turnover dynamics of retinal macrophages at the steady state and after insult.

In conclusion, our study, providing an atlas of non-neuronal cells in the adult mouse retina, identifies key events along the path of neuronal degeneration. Many features of this response also recur in other CNS pathological settings, beyond the ONC model and the retina, thus offering a resource applicable for studying neuroimmune processes at large. The analysis of concerted multicellular events underlying tissue biology holds promise for making headway in CNS repair.

## Acknowledgements

We thank I. Avraham-Davidi, T. van Zyl, R. Kedmi, I. Shachar and M. Schwartz for helpful discussions; M. Schwartz and M. Mack for kindly providing the MC-21 antibody; Y. Okunuki and K. Connor for their help with the PLX experiment; L. Jerby-Arnon for assistance with DIALOGUE analysis, K. Dey and K. Jagadeesh for help with GWAS analysis, and R. Harpaz for help with RPE image analysis; Z. Niziolek, S. Turney, Y. Li, R. Schaffer, M. Laboulaye, E. Martersteck, T. Delorey, D. Phillips, and staff members of the Harvard University Bauer Core Facility and the Koch Institute Histology Core for technical assistance. We thank L. Gaffney and A. Hupalowska for help with figure preparation, and the Richter family for support. This study was supported by the Human Frontier Science Program, the Center for Integration in Science of the Israeli Ministry of Absorption, HHMI, the Klarman Family Foundation, Wings for Life Spinal Cord Research Foundation, and NIH grants EY028633 and MH105960. The funders had no role in the study design, experiments performed, data collection, data analysis and interpretation, or preparation of the manuscript.

## Author Contributions

I.B. and A.R. conceived the study. I.B., I.E.W., K.S., J.R.S. and A.R. designed experiments and analyzed data. I.B., I.E.W., A.J., M.S., G.B., N.M.T. and W.C. performed experiments. J.D., W.Y. and K.S. developed computational approaches and analyzed scRNA-seq data. Z.H., J.R.S. and A.R. provided supervision and acquired funding. I.B., J.D. and A.R. wrote the manuscript, with input from all authors.

## Methods

### Mice

All animal experiments were approved by the Institutional Animal Care and Use Committees (IACUC) at Harvard University and Children’s Hospital, Boston. Mice were maintained in pathogen-free facilities under standard housing conditions with continuous access to food and water. All experiments were carried out in adult mice from 6 to 16 weeks of age and included both males and females. The following mouse strains were used: C57BL/6J (JAX #000664), Vglut2-ires-cre (*Slc17a6^tm2(cre)Lowl^/*J (Vong et al., 2011)) crossed to the cre-dependent reporter Thy1-stop-YFP Line#15 (B6.Cg-Tg(Thy1-EYFP)15Jrs/J (Buffelli et al., 2003), and CCR2^RFP^ (B6.129(Cg)-*Ccr2^tm2.1Ifc^*/J; JAX #017586) (Saederup et al., 2010) crossed to C57BL/6J mice to generate heterozygotes.

### Optic Nerve Crush (ONC)

Optic nerve crush (ONC) injury was performed as previously described (Benhar et al., 2016; Tran et al., 2019). Briefly, after anesthesia with ketamine/xylazine (ketamine 100-120 mg/kg and xylazine 10 mg/kg), the optic nerve was exposed intraorbitally and crushed with fine forceps (Dumont #5 FST) for 2 s approximately 0.5-1 mm behind the optic disc. All surgeries were performed by an experienced surgeon, who visually confirmed optic nerve crush during the procedure.

### Cell preparation and flow cytometry

Mice were intracardially perfused with PBS. Upon dissection, eyes were visually inspected for damage or blood, which were used as criteria for exclusion. For scRNA-seq experiments, retinas or RPE from 2-8 mice were typically pooled per condition.

*Retina:* Eyes were dissected in AMES solution (equilibrated with 95% O_2_/5% CO_2_). Retinas, separated from eyecup and ciliary body, were digested in papain, and dissociated to single cell suspensions using manual trituration in ovomucoid solution. Cells were spun down at 450 g for 8 minutes and resuspended in AMES+4%BSA to a concentration of 10 million cells per 100 ml.

*RPE:* RPE sheets were isolated from posterior eyecups based on a protocol adapted from Fernandez-Godino et al. (Fernandez-Godino et al., 2016). Briefly, eyecups were dissected in Hank’s Balanced Salt Solution (HBSS)^-^+HEPES and incubated in 0.25% trypsin/EDTA for 45 minutes at 37°C, after which RPE sheets were released by manual shaking into 20% fetal bovine serum (FBS) in HBSS^+^-HEPES, spun down at 240 g for 2 minutes and incubated in 0.05% Trypsin/EDTA for 10 minutes at 37°C. Cells were washed, resuspended and triturated 50 times in 5% FBS/HBSS^+^-HEPES and filtered for staining and cell sorting.

The following antibodies were used, conjugated to various fluorophores: anti-CD90.2 (Thy1.2, clone 53-2.1, ThermoFisher Scientific), CD45 (clone 30-F11, BD Pharmingen), CD11b (clone M1/70, BioLegend), CD115 (clone AFS98, BioLegend), Ly6C (clone HK1.4, BioLegend), CD140a (clone APA5, BioLegend), GLAST (clone ACSA-1, Miltenyi Biotec). Cells were incubated with antibodies for 15 minutes, washed with an excess of media, spun down and resuspended again in AMES+4%BSA at a concentration of 7 million cells/ml. The live cell marker Calcein Blue (ThermoFisher Scientific) was added just prior to FACS. Cellular debris, doublets, and dead cells were excluded. For scRNA-seq, cells were gated on CD45, GLAST (EAAT1, *Slc1a3*) and CD140a (PDGFRɑ) to enrich for immune cells, Müller glia and astrocytes, respectively. Since GLAST is expressed by both Müller glia and astrocytes, and the former are much more abundant, astrocytes were specifically enriched using the marker CD140a (Pdgfra), which is expressed by retinal astrocytes (Macosko et al., 2015; Takahama et al., 2017). Cells were collected into 100μl of AMES+4%BSA per 25,000 sorted cells. Following collection, cells were spun down and resuspended in PBS+0.1%non-acetylated BSA at a concentration range of 500-2000 cells/μl for droplet-based scRNA-seq per manufacturer’s instructions (10x Genomics).

### 3’ droplet-based scRNA-seq

Single cell libraries were prepared using the Single-cell gene expression 3’ v2 or v3 kit on the Chromium platform (10X Genomics, Pleasanton, CA) following the manufacturer’s protocol. On average, approximately 7,000-12,000 single cells were loaded on each channel and approximately 3,000-7,000 cells were recovered. Quantification and quality control analyses were performed using the Agilent Bioanalyzer High Sensitivity DNA assay. Libraries were sequenced on Illumina NextSeq 500 or HiSeq X platforms (Paired end reads: Read 1, 26 bases; Read 2, 55 bases for NextSeq, 98 bases for HiSeq). From the retina, 2-5 independent experiments per time point were performed for scRNA-seq. From the eyecup, one experiment was performed for each time point.

### Histological methods and imaging

Eyes were collected from mice intracardially perfused with 4% paraformaldehyde (PFA), and post-fixed in 4% PFA for an additional 15-30 minutes or 1 hour for RPE whole-mounts. Eyes were transferred into PBS until dissection, following which retinas or eyecups were either used for whole-mount IHC or posterior eyecups were sunk in 30% sucrose and embedded in tissue freezing media (OCT) to cryosection into 20-25μm-thick cross-sections. For formalin-fixed paraffin-embedded (FFPE) sections, eyes from mice intracardially perfused with PBS were post-fixed in 10% formaldehyde at room temperature overnight and transferred to 70% ethanol prior to paraffin embedding and sectioning into 5μm-thick sections.

To immunostain retinal/RPE whole-mounts, retinas or posterior eyecups with the retina removed were incubated in protein block (5% normal serum, 0.3% triton-x, 1x PBS) for 3 hours at room temperature or overnight at 4°C, followed by incubation with primary antibodies (in protein block) for 3-7 days, and secondary antibodies (in 1x PBS) overnight. All antibody incubations were done at 4°C with gentle rocking.

For IHC on cryosections, slides were incubated for 1h in protein block, primary antibody incubation overnight, and secondary antibodies for 2-3h. Initial block and secondary antibody incubation were done at room temperature and primary antibody incubation at 4°C.

For IHC on FFPE sections, slides were baked at 60°C for 30 minutes, followed by deparaffinization and rehydration through a series of successive incubations in Xylenes, ethanol (100%, 95%, 70%, 50%) and finally distilled water. Antigen retrieval was performed with Antigen Unmasking Solution (Vector Laboratories) in a vegetable steamer. Sections were blocked in CAS-Block™ Histochemical Reagent (ThermoFisher Scientific) for 30 minutes at room temperature and incubated overnight with primary antibodies in CAS block at 4°C in a humidity chamber.

After washes in PBS, sections were incubated with secondary antibodies in PBS for 1 hour at room temperature, before mounting with DAPI.

Fluorescence *in situ* hybridization (FISH) was performed on FFPE sections using the commercially available RNAscope Fluorescent Multiplex Assay (ACD), or on whole-mount retina with HCR (Molecular Instruments), according to manufacturers’ instructions.

Zeiss LSM 710 and Olympus FV-1000 confocal microscopes were used for imaging. Images were merged, cropped and optimized using ImageJ (Fiji) (Schindelin et al., 2012) and Adobe Photoshop, and arranged using Adobe Illustrator.

### Antibody treatment for monocyte depletion

CCR2^+^ monocytes were depleted by intraperitoneal injections of anti-CCR2 monoclonal antibody (MC-21 (Mack et al., 2001); 400μg), starting one day before optic nerve crush and every other day until sacrifice (up to 5 injections). Depletion efficiency was confirmed in the blood using flow cytometry.

### PLX5622 treatment for myeloid cell depletion

Mice were fed with control chow (AIN-76) or with chow containing 1,200 ppm of the Csf1r antagonist PLX5622 (Plexxikon, Inc.), as previously described (Okunuki et al., 2019) starting 1 week before ONC and for an additional 1 week until sacrifice.

### ScRNA-seq pre-processing

CellRanger 2.1.0 (10X Genomics) was used for read demultiplexing, alignment to the mouse genome and unique molecular identifier (UMI) counting and collapsing. The top 6,000 cells from each experiment with the largest number of UMI were used for further analysis (Tran et al., 2019). In total, 121,309 cells expressing at least 400 genes per cell were retained for downstream analysis.

### Batch correction and clustering: Retina

scPhere (Ding and Regev, 2021), a deep learning-based method, was used to merge all 121,309 cells from different time points by taking time after crush and mouse strain as the batch vectors. For scPhere analysis, a latent space of dimension 10 was used so each cell was mapped to a 10-sphere. The latent representation of cells were clustered using the Louvain community detection algorithm (Blondel et al., 2008; Levine et al., 2015) to produce 45 clusters. These clusters were assigned to 25 putative cell types/groups (including cell doublets and low-quality cell clusters) using a previously-described automatic cell type assignment approach (Ding et al., 2020) followed by manual inspection (**Fig. 1b,c**). Cell clusters that expressed neuronal markers (‘neurons’), expressed markers genes of two cell types (‘contaminants’), or had very small numbers of detected genes per cell and did not express cell subtype-specific marker genes (‘Low UMI’) were discarded, retaining 107,067 cells assigned to 14 broad groups for further analysis. These 107,067 cells were partitioned into three categories by cell class: astrocytes (18,959 cells), Müller glia (54,565 cells), and all other cells (33,543 cells, predominantly immune cells). (As each cell category was separately enriched and sorted; we only calculate proportions between subsets within a category, but not across the categories.) To identify cell subtypes and very rare cell types, the cells from each compartment were then analyzed separately, either by re-clustering these cells using the Louvain community detection algorithm (Blondel et al., 2008; Levine et al., 2015) implemented in Seurat (Hao et al., 2021), specifically using scPhere low-dimensional representations (astrocytes and Müller glia), or by rerunning scPhere for batch-correction and dimensionality reduction, then re-clustering scPhere results (immune cells). This analysis further identified three Müller glia subsets (**Fig. 3d**), five astrocyte subsets (**Fig. 3g**), and 22 immune cell subsets, as well as removed another 3,116 ‘contaminant’ cell profiles from the immune cell compartment, retaining 30,427 immune cells in total (including some stromal cell subsets) after filtering (**Fig. 2a**).

### Batch correction and clustering: Eyecup

The data was analyzed using R package Seurat following previously described methods (Jacobi et al., 2022). Briefly, cells collected at each time point were identified as one batch and the function “sctransform” was applied to each batch for normalization. The top 3,000 features were used for integration of all batches using function “IntegrateData”. Markers for each cluster were identified using the function “FindAllMarkers” with the “Mast” algorithm. Cell types were annotated based on canonical markers. A low quality RPE cluster was filtered post hoc based on high expression of mitochondrial genes together with a low number of genes per cell relative to the other RPE clusters.

### Differential expression analysis

Logistic regression was used for differential gene expression analysis, taking both the log2 transformed total number of detected genes in each cell and the mouse strain as covariates, and the Seurat implementation for differential expression analysis (Hao et al., 2021).

### Gene Ontology enrichment analysis

To identify biological processes enriched in RPE clusters, we tested for enrichment in Gene Ontology (GO) biological process gene sets from the Molecular Signatures Database (MSigDB) (Liberzon et al., 2015; Subramanian et al., 2005), which uses the hypergeometric distribution.

### Gene signature analysis

Gene signature scores were calculated as previously described (Tirosh et al., 2016), with the implementation in the Seurat package (Hao et al., 2021). The interferon gene signature was obtained from ImmGene (Mostafavi et al., 2016).

### MC-21 (monocyte depletion) data analysis

To help annotate cells from MC-21 antibody treated mice and control mice, cell profiles were coembedded using scPhere (Ding and Regev, 2021) with all immune cell profiles (**Fig. 2a**). A *k*-nearest neighbor classifier (*k*=15) trained on the immune cells with cell state annotations (**Fig. 2a**) was used to annotate cells from both control and MC-21-treated mice. Prior to this analysis, likely non-immune cell contaminants were first separately removed from the profiled cells from both control and MC-21-treated mice. Specifically, to remove these non-immune cells, the full analysis was performed with scPhere for batch correction and dimensionality reduction, taking experiment as batch vector, and then scPhere’s low-dimensional representations of cell profiles (in 5 dimensions) were clustered using the Louvain algorithm (Blondel et al., 2008; Levine et al., 2015) as implemented in Seurat (Hao et al., 2021). Non-immune cells were manually annotated and removed before further analysis.

### Optimal transport and RNA velocity analysis

The Waddington-OT package (Schiebinger et al., 2019) was used with default parameters as in the Waddington-OT online tutorial (https://broadinstitute.github.io/wot/tutorial/) to trace the likely ancestors of cells from two subsets of *Ms4a7*^+^ macrophages.

For myeloid cell differentiation trajectory analysis, velocyto (La Manno et al., 2018) and scVelo (Bergen et al., 2020) were used. Specifically, after running CellRanger, velocyto was used to calculate the intronic and exonic UMIs for each cell passing the initial filtering, and then scVelo was used to calculate the likely differentiation potential for each cell. Although scVelo can be impacted by batch effects (*e.g.*, samples confounded with time), there was no apparent strong batch effect between monocytes, macrophages, or dendritic cells. Thus, scVelo was run with velocyto results as inputs.

### Identification of multicellular programs

The DIALOGUE method was used to identify multicellular programs (Jerby-Arnon and Regev, 2022) only with resident cells (astrocytes, Müller glia and microglia), without partitioning them into more granular cell states. We followed the default settings for DIALOGUE analysis. The number of multicellular programs (MCP) *k* was set to 6. We only considered ‘ctrl’ or ‘crush’ and tested the association of this binary feature with MCPs (DIALOGUE only supports binary feature for the current version -- the ‘pheno’ parameter). DIALOGUE also takes batch vectors as covariant in analysis and we used mouse strain as the ‘conf’ parameter. DIALOGUE needs the full non-sparse matrix of each compartment as an input. Thus, for each compartment, we randomly subsampled 20,000 cells for analysis if the total number of cells from that compartment was greater than 20,000.

### Receptor-ligand based interaction analysis

A set of 2,392 manually-curated ligand-receptor pairs (**Supp. Table 3**), obtained from recent publications (Baccin et al., 2020; Hou et al., 2020), and manually updated (removing pairs that didn’t have mouse homologs or convincing literature support in mice, correcting homologs, ultimately adding 13 pairs), was used to identify putative cell-cell interactions between cell types. There are several challenges for current methods for inferring the cell-cell interactions from scRNA-seq data. First, cell types expressing receptors or ligands whose cognate partner is highly expressed ubiquitously across all cells, can become interaction hubs. Second, some methods consider only pairwise interactions but not multi-subunit complexes. Third, because of measurement errors (*e.g.*, ambient RNAs), ligand or receptor expression can be detected at least at low levels in cell types that do not actually express these genes, and will add noise to the interaction strength ranks between cell types. This is likely given the large number of ligand-receptor pairs scored (above 2,000).

To address these problems, a *k*-nearest neighbor approach was used to estimate the interaction strength between cell types. Specifically, for a gene *g* of a cell type, its score *s_g_* is defined as the total number of detected UMIs of gene *g* in that cell type divided by the total number of UMIs of gene *g* across all cell types. If *g* is a member of a multi-subunit complex with *m* subunits, the weight for that complex is defined as the geometric mean of scores of all members. Then the final interaction score between a ligand-receptor pair is the product of the weights of the two interaction complexes. For any two cell types *i* and *j*, the interaction score is calculated for each ligand-receptor pair. Finally, the interaction strength between two cell types is the sum of the top *k* interaction scores. These interaction strength scores are used to rank the interaction partners of a cell type. Only used the top *k* interaction pairs were used to decrease the influence of the background interaction noise.

### Mapping disease genes

The retinal disease gene lists were generated from the following sources: POAG (Gharahkhani et al., 2021); VDD and VCDR (Han et al., 2021); all other diseases from the Retinal Information Network (RetNet, https://sph.uth.edu/retnet/), downloaded on 12/16/2021. Analysis was performed as previously described (van Zyl et al., 2022). Briefly, the retina and eyecup datasets were merged and re-normalized before plotting. Genes from each disease catalog were filtered to retain only those expressed in more than 10% of cells. Values were z-scaled in each row of the heat map and genes were ordered by the peak values in the columns.

---

**Supp. Fig. 1:**
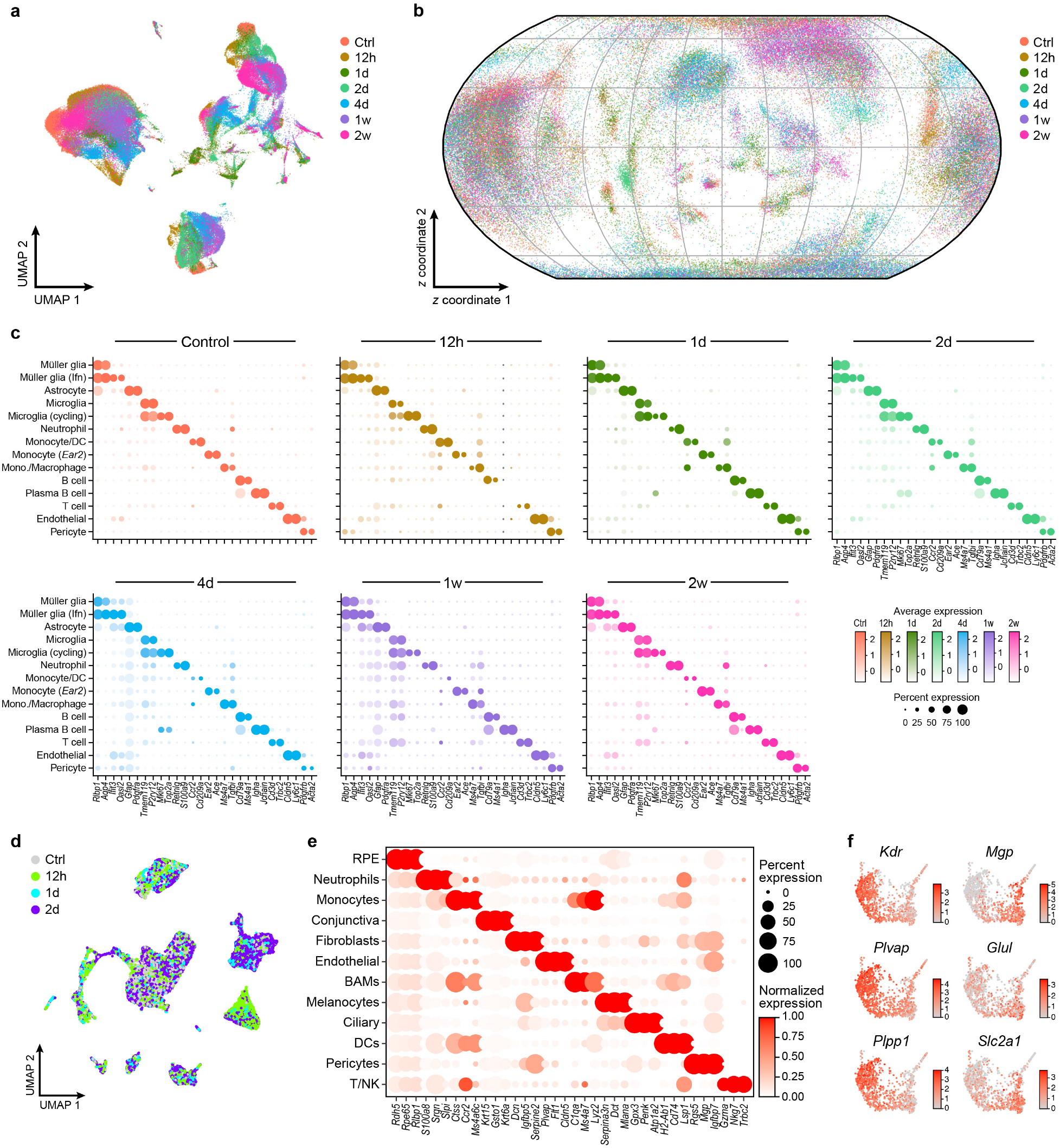
Changes in cell composition and stability in cell marker expression throughout the time course. **a,** 2D Uniform Manifold Approximation and Projection (UMAP) for 121,309 single cells profiled from the retina across the time course, colored by time point. **b,** ScPhere embedding of 121,309 single cell profiles (dots) from the retina, projected to 2D by the Equal Earth map projection method, colored by time point (legend). **c,** Retinal cell type marker gene expression is stable across the time course. Shown are fraction of expressing cells (dot size) and mean expression levels in expressing cells (dot color) of selected marker genes (columns) across 14 non-neuronal cell types (rows), plotted at each time point. **d,** UMAP for 21,275 cells profiled from mouse posterior eyecup, colored by time point. **e,** Fraction of expressing cells (dot size) and normalized expression in expressing cells (dot color) of selected marker genes (columns) across 12 identified cell types in the eyecup (rows). **f,** Endothelial cells collected from mouse eyecup, colored by expression of genes associated with choroidal Kdr^hi^ (left) and Kdr^lo^ (right) cells (Lehmann et al., 2020).

**Supp. Fig. 2:**
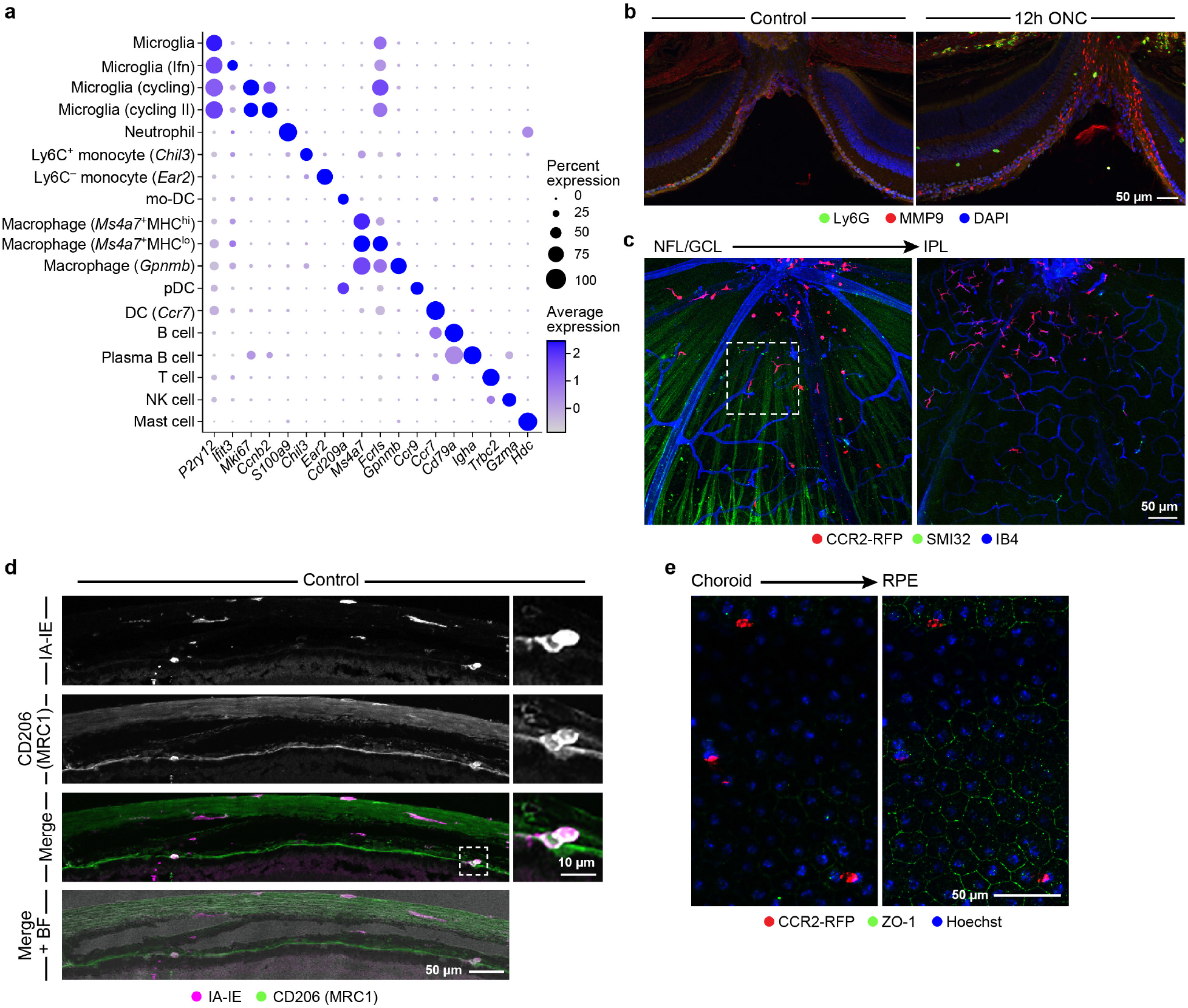
Visualizing immune cells in the retina and eyecup. **a,** Immune cell subsets and their markers. Fraction of expressing cells (dot size) and mean expression levels in expressing cells (dot color) of selected marker genes (columns) across 18 identified immune cell subsets (rows). **b,c,** Representative images of IHC on retinal sections (**b**) showing that neutrophils (LY6G^+^, green) are increased 12h after ONC, express MMP9 (red) and localize proximal to the optic nerve head (ONH) (n = 3 per time point), and on retinal whole-mounts from CCR2^RFP/+^ mice at 2dpc (**c**) showing CCR2-RFP^+^ cells (red) at the levels of the GCL and IPL. Higher magnification inset of the outlined region in **c** is shown in Fig. 2d (n = 2). **d,e,** Myeloid cells in the uninjured eyecup. **d,** Resident macrophages in the eyecup express IA-IE and CD206. Representative image of IHC on eyecup sections for IA-IE (magenta) and CD206 (green). Inset shows double positive cells. BF, brightfield (n = 2-3, representative of two independent experiments). **e,** Representative images of IHC on eyecup whole-mounts from CCR2^RFP/+^ mice showing that the majority of CCR2-RFP cells (red) in naïve eyecup are located posterior to the RPE, at the level of the choroid. ZO-1 (green) depicts tight junctions between individual RPE cells and nuclei are stained with Hoechst (blue) (n = 2, representative of three independent experiments).

**Supp. Fig. 3:**
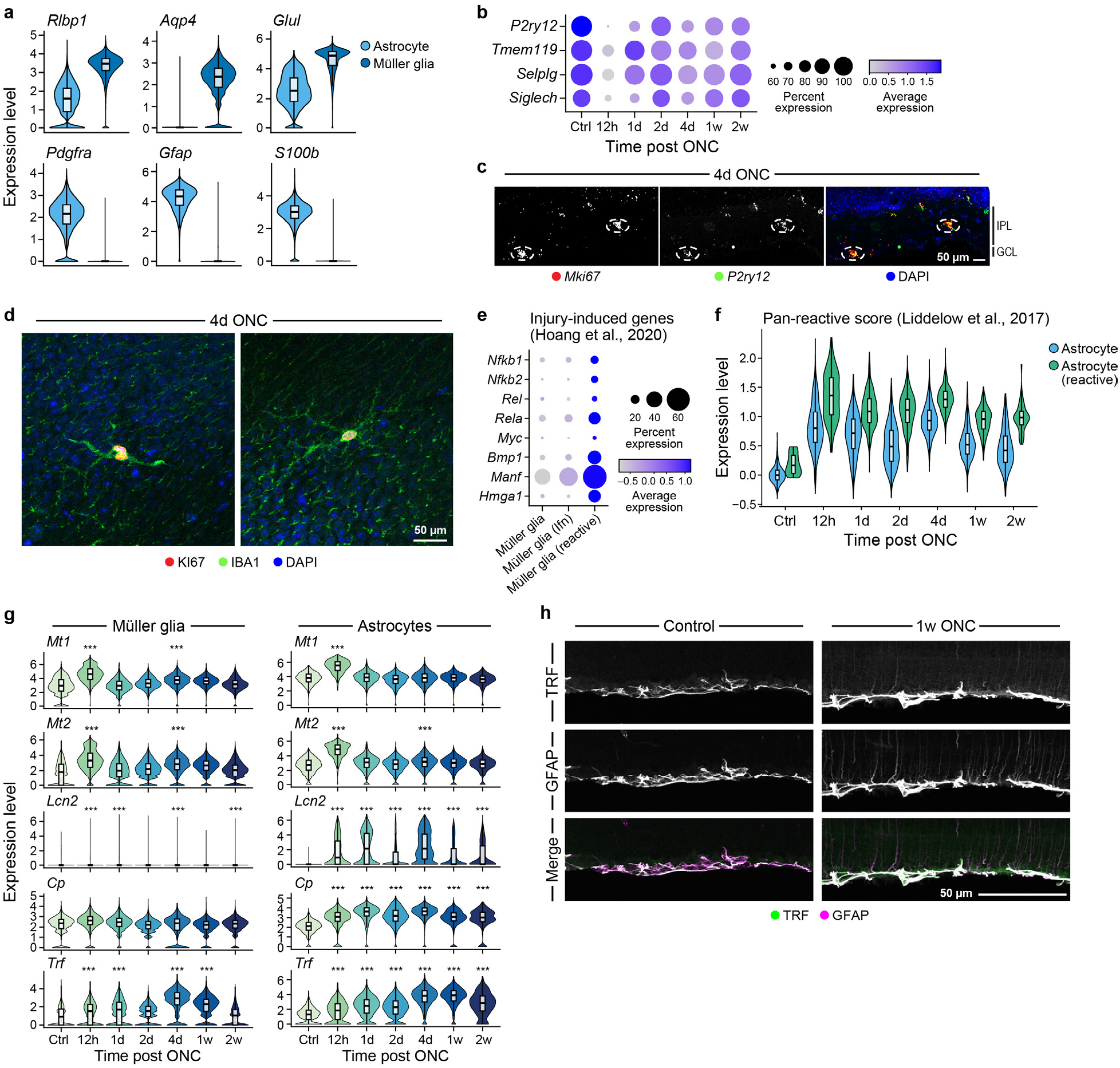
Glial reactivation and microglial proliferation after ONC. **a,** Distribution of expression of genes differentially expressed between Müller glia and astrocytes. **b,** Fraction of expressing microglia cells (dot size) and mean expression levels in expressing cells (dot color) of microglia signature genes across time, showing decreased expression after ONC. **c,d,** Microglia proliferate after ONC. Shown are representative images from 4dpc of smFISH on retinal sections showing *P2ry12^+^* microglia (green) expressing the proliferation marker, *Mki67* (red) (**c**) (n = 3), and of whole-mounts showing IBA1^+^ microglia (green) expressing KI67 protein (red) (**d**) (n = 3). **e,** Fraction of cells (dot size) in each Müller glia subset, and mean expression level in expressing cells (dot color) of injury-induced genes described by Hoang et al. (Hoang et al., 2020). **f,** Distribution of expression scores for signature genes of pan-reactive astrocytes (Liddelow et al., 2017) across astrocyte subsets throughout the time course. **g,h,** Glial reactivity includes upregulation of metal metabolism gene expression. **g,** Distribution of expression of metal metabolism genes by Müller glia (left) and astrocytes (right) across the time course. \*\*\**p* < 0.001, fold change > 1.5 relative to ctrl. **h,** Representative images of IHC on sections from control (left) and 1wpc (right) retina showing expression in astrocytes and Müller glia (GFAP, magenta) of transferrin (TRF, green), which is increased after ONC (n = 3 per time point).

**Supp. Fig. 4:**
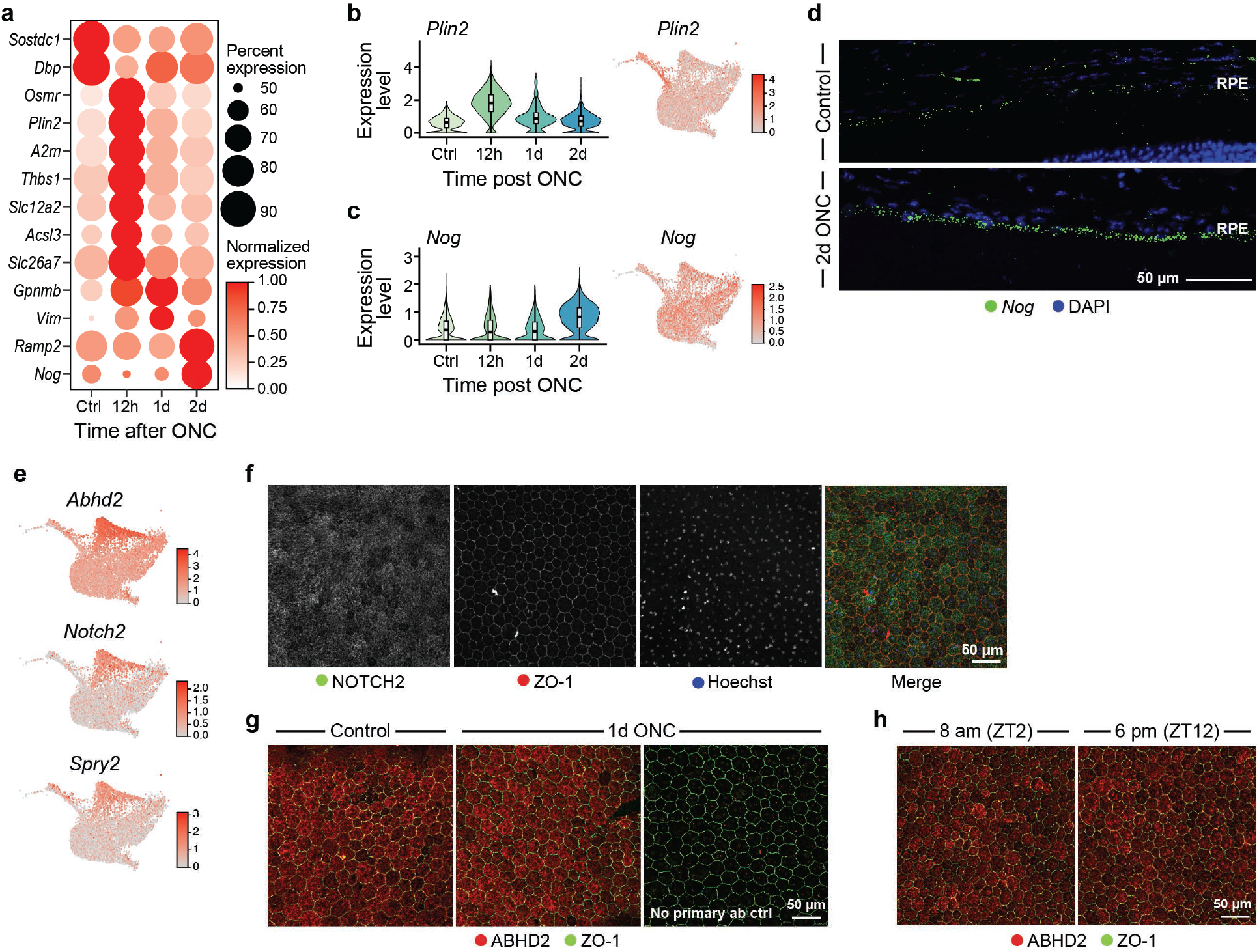
Baseline and ONC-induced heterogeneity in the RPE. **a-d,** ONC-induced changes in the RPE. **a,** Fraction of expressing cells (dot size) and normalized expression level in expressing cells (dot color) of differentially expressed genes (rows) in the RPE between different time points (columns). **b,c,** Distribution of expression of *Plin2* (**b**) and *Nog* (**c**) on RPE cells by time (left) and across subsets (right). **d,** Representative image of smFISH for *Nog* (green) on uninjured (Control) and 2dpc RPE (n = 3 per time point). **e-h,** Identification of heterogeneity in the RPE, which is maintained regardless of injury or circadian cycle. **e,** RPE cells colored by expression of C4 top DEGs. **f,** Representative images of IHC on RPE whole-mounts showing heterogeneity in NOTCH2 protein expression (green). ZO-1 (red) depicts tight junctions between individual RPE cells and nuclei are stained with Hoechst (blue) (representative of five independent experiments with n = 2-4). **g,h,** Representative images of IHC on RPE whole-mounts for ABHD2 (red) expression before and after ONC (n = 3, representative of four independent experiments) (**g**) and at two distinct time points in the circadian cycle (**h**). (ZT=Zeitgeber Time, indicates lights on; ZT2 is 2 hours after lights on and ZT12 is 2 hours before lights off) (n = 3 per time point). The panel on the right in (G) shows a negative control in which the primary antibody was omitted to show staining specificity.

**Supp. Fig. 5:**
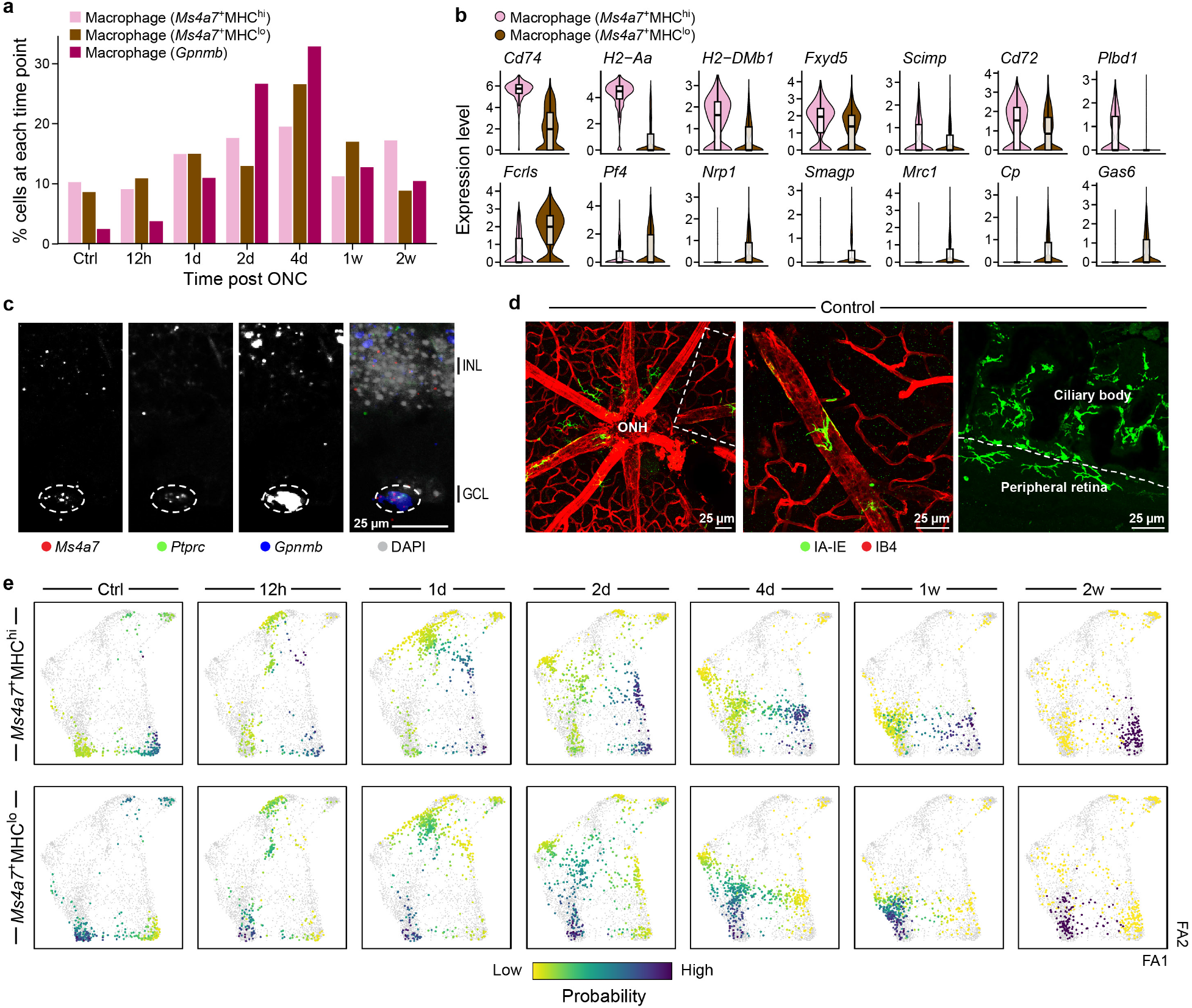
Dynamics of resident and infiltrating mononuclear phagocytes in the retina. **a,** Changes in frequency (y axis) of *Ms4a7*^+^MHC^hi^, *Ms4a7*^+^MHC^lo^ and *Gpnmb* macrophages across the time course (x axis). **b,** Distribution of expression of genes associated with MHC-II^hi^ and MHC-II^lo^ BAMs (Van Hove et al., 2019) on retinal *Ms4a7*^+^MHC^hi^ and *Ms4a7*^+^MHC^lo^ macrophages. **c,** Representative image of smFISH on retinal sections showing that *Gpnmb*^+^ (blue) immune cells (*Ptprc*, green) are also *Ms4a7*^+^ (red) (n = 3). **d,** Representative images of IHC on uninjured (Control) retinal whole-mounts showing perivascular and CB-adjacent IA-IE^+^ (green) cells. Blood vessels are labeled with IB4 (red) (representative of five independent experiments with n = 1-3). The middle panel shows a zoomed-in image of the perivascular macrophage in the inset of the left image. **e,** Force-directed layout view of monocyte and macrophage subsets with optimal transport analysis. Each dot is a cell, color-coded by ancestor probabilities, as estimated by Waddington-OT (Schiebinger et al., 2019).

**Supp. Fig. 6:**
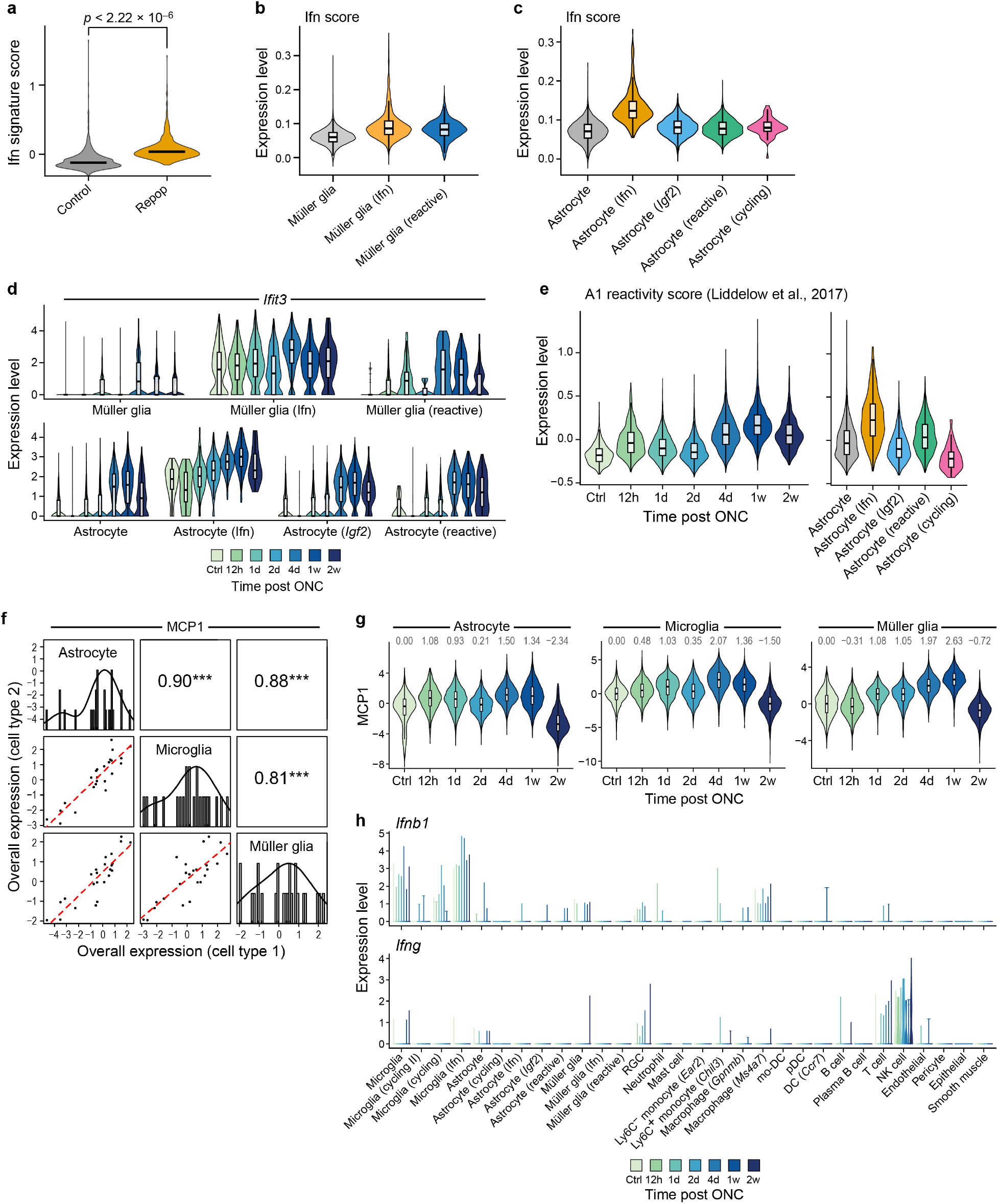
Rare Ifn-responding glial subsets and a co-regulated Ifn-response program in the injured retina. **a,** Distribution of expression of genes differentially expressed in our Ifn microglia by microglia repopulating the brain after depletion (Repop) compared to control microglia (Huang et al., 2018a). **b,c,** Distribution of expression scores of a previously curated (Mostafavi et al., 2016) Ifn gene signature across Müller glia (**b**) and astrocyte (**c**) subsets. **d,** Distribution of *Ifit3* expression across time in Müller glia (top) and astrocytes (bottom) across subsets. **e,** Distribution of expression scores for signature genes of “A1” astrocytes (Liddelow et al., 2017) across astrocytes throughout the time course (left) and by subset (right). **f,g** The Ifn response is part of a coordinated multicellular program increased after ONC. **f,** Off diagonal panels: Comparison of overall expression scores (y and x axes) for each cell component of MCP1 (rows and columns, labels on diagonal) across the samples; lines correspond to linear fit. Pearson correlation (r) and significance (\*\*\**p* < 0.001) are shown in the panels above the diagonal. Diagonal panels: Distribution of overall expression scores for each cell type component, with kernel density estimates (Jerby-Arnon and Regev, 2022). **g,** Distribution of MCP1 expression scores by astrocytes, microglia and Müller glia along the time course. Numbers above violins indicate median expression scores. **h,** Distribution of expression of *Ifnb1* (top) and *Ifng* (bottom) by cell types in the retina across time (x axis).

**Supp. Fig. 7:**
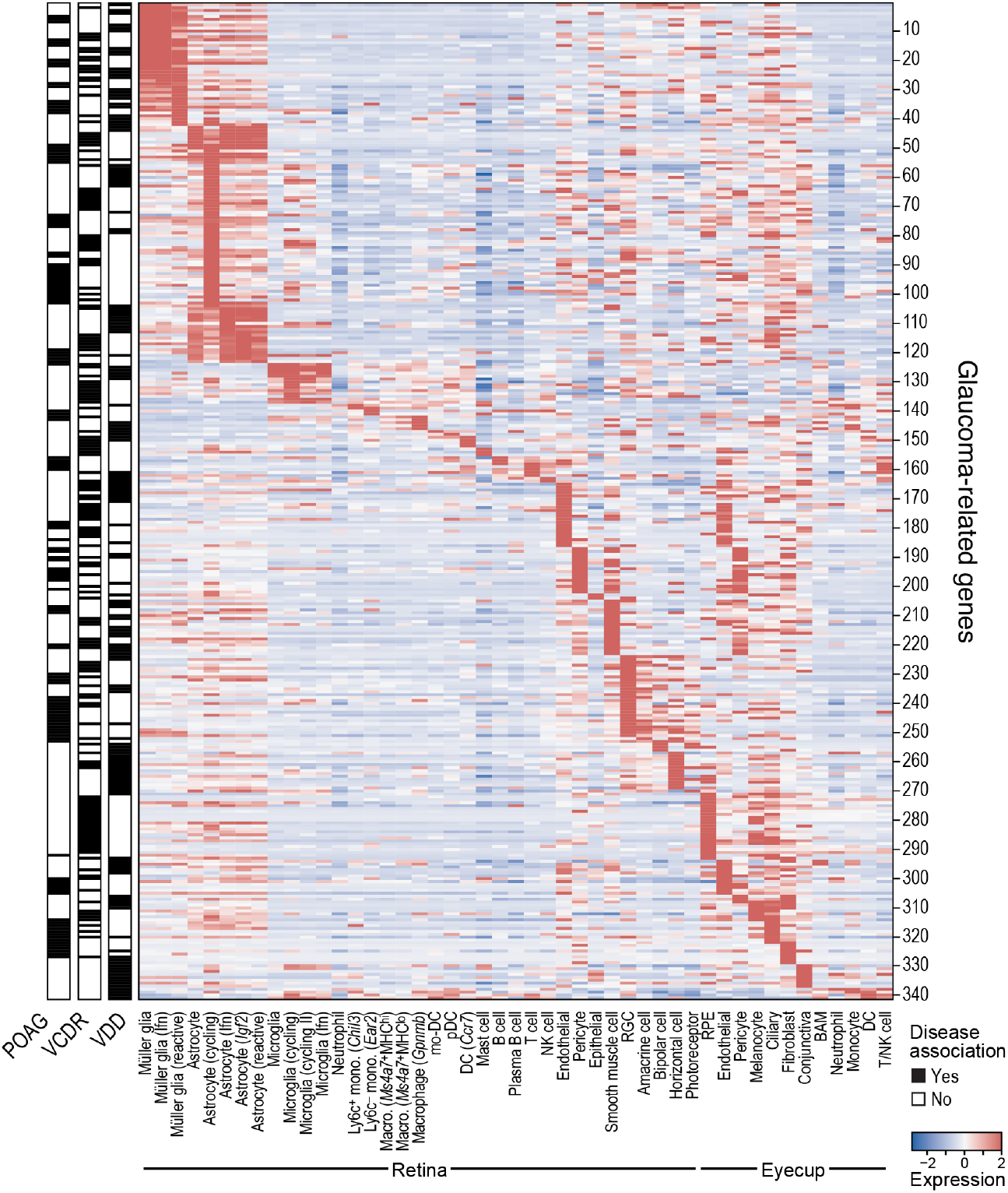
Expression of glaucoma related genes in mouse retinal and eyecup cell types. Genes implicated in glaucoma are enriched in retinal glia. Average expression (z-score, red/blue bar) of genes associated with primary open angle glaucoma (POAG), vertical-cup-to-disc ratio (VCDR) and vertical disc diameter (VDD) (rows; black bars on left; **Supp. Table 2**) across cell types in the retina and eyecup (columns).

**Supp. Fig. 8:**
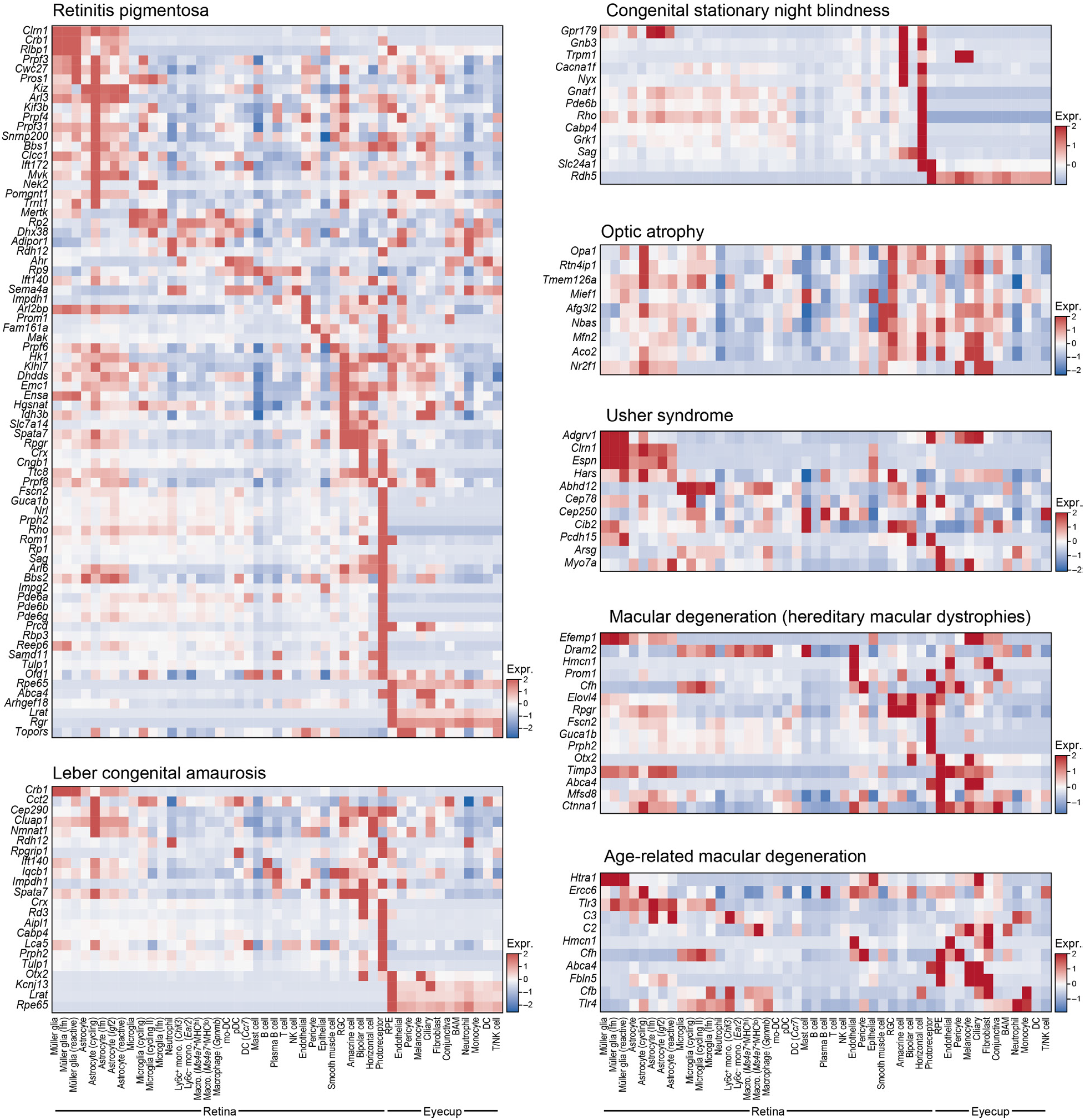
Expression of retinal disease related genes in retinal and RPE cell types. Average expression (z-score, red/blue bar) of genes (rows) implicated in each of seven groups of blinding diseases, across cell types in the retina and eyecup (columns).

## Table Legends

**Supp. Table 1.** MCP1 up and downregulated genes in astrocytes, microglia and Müller glia.

**Supp. Table 2.** Genes associated with POAG, VCDR and DiscD. Index numbers correspond to the row numbers in Supp. Figure 7.

**Supp. Table 3.** Ligand-receptor list used for inferring cell-cell interactions, based on the lists from Baccin et al., 2020; Hou et al., 2020.

**Supp. Table 4.** List of antibodies used in this study.

